# A FACT-ETS-1 Antiviral Response Pathway Restricts Viral Replication and is Countered by Poxvirus A51R Proteins

**DOI:** 10.1101/2023.02.08.527673

**Authors:** Emily A. Rex, Dahee Seo, Sruthi Chappidi, Chelsea Pinkham, Sabrynna Brito Oliveira, Aaron Embry, David Heisler, Yang Liu, Karolin Luger, Neal M. Alto, Flávio Guimarães da Fonseca, Robert Orchard, Dustin Hancks, Don B. Gammon

## Abstract

The FACT complex is an ancient chromatin remodeling factor comprised of Spt16 and SSRP1 subunits that regulates specific eukaryotic gene expression programs. However, whether FACT regulates host immune responses to infection was unclear. Here, we identify an antiviral pathway mediated by FACT, distinct from the interferon response, that restricts poxvirus replication. We show that early viral gene expression triggers nuclear accumulation of specialized, SUMOylated Spt16 subunits of FACT required for expression of ETS-1, a downstream transcription factor that activates a virus restriction program. However, poxvirus-encoded A51R proteins block ETS-1 expression by outcompeting SSRP1 for binding to SUMOylated Spt16 in the cytosol and by tethering SUMOylated Spt16 to microtubules. Moreover, we show that A51R antagonism of FACT enhances both poxvirus replication in human cells and viral virulence in mice. Finally, we demonstrate that FACT also restricts unrelated RNA viruses, suggesting a broad role for FACT in antiviral immunity. Our study reveals the <u>F</u>ACT-<u>E</u>TS-1 <u>A</u>ntiviral <u>R</u>esponse (FEAR) pathway to be critical for eukaryotic antiviral immunity and describes a unique mechanism of viral immune evasion.

## Main

Activation of antiviral gene expression after viral infection is a critical aspect of host antiviral responses. Key among these responses in mammals is the Type I Interferon (IFN) response, which induces hundreds of antiviral genes during infection^1, 2^. The importance of the IFN response is underscored by the fact that virtually all mammalian viruses encode IFN antagonists^1, 2^. However, relatively little is known regarding antiviral transcriptional responses that evolved prior to vertebrate-specific IFN responses.

Poxviruses are large, cytoplasmic DNA viruses that infect animals and humans worldwide. For example, variola virus (VARV), the causative agent of smallpox, was responsible for one of the deadliest infectious diseases in human history^3^. Despite the successful eradication of smallpox by 1979, zoonotic poxvirus infections remain a major public health concern. This is underscored by the declaration by the World Health Organization that the 2022-2023 outbreak of Mpox (formerly known as “monkeypox”) is a public health emergency of international concern^4, 5^. Poxvirus infections are often highly pathogenic and associated with severe host immune suppression due to the action of poxvirus-encoded antagonists that inhibit various host immune responses^6^. Notably, the identification and characterization of poxviral immune evasion factors has often revealed new functional insights into the host factors and pathways they target^7^.

Previously, we showed the early A51R protein encoded by the mammalian poxvirus, vaccinia virus (VV), to be required for robust VV replication in mammalian cells and for virulence in mice^8^. Although A51R deficiency did not alter VV susceptibility to IFN treatment in mammalian cell cultures, curiously, A51R expression alone could promote RNA virus replication in non-permissive insect cells^8^. These observations suggested that A51R inhibits undefined eukaryotic antiviral responses that arose prior to the IFN response during evolution.

The human “<u>Fa</u>cilitates <u>C</u>hromatin <u>T</u>ranscription” (FACT) complex is an evolutionarily-conserved, chromatin remodeler that requires interaction between human Suppressor of Ty 16 homolog (hSpt16) and Structure-Specific Recognition Protein-1 (SSRP1) subunits to function. FACT regulates mRNA transcription by controlling chromatin accessibility to transcriptional machinery^9, 10, 11^, but is not required for all mRNA transcription and instead localizes to discrete genomic loci to regulate specific cellular genes^12–14^. However, the biological relevance of FACT-mediated gene expression programs to immunity is largely unknown.

Here, we reveal poxvirus A51R proteins to antagonize a novel, FACT-mediated antiviral pathway that is activated by poxvirus early gene expression. We demonstrate that VV A51R competes with SSRP1 to specifically, and directly, bind a novel SUMOylated form of hSpt16 in the cytoplasm of infected cells. This interaction tethers SUMOylated hSpt16 to microtubules (MTs), preventing its nuclear accumulation and activation of a FACT-dependent antiviral gene expression program distinct from the IFN response. Moreover, we show A51R proteins from multiple poxviruses specifically bind SUMOylated hSpt16, suggesting FACT antagonism is a conserved function of A51R proteins. We demonstrate that hSpt16 SUMOylation promotes hSpt16 binding to monoubiquitinated H2B histone (which marks active transcription sites in chromatin^15^) during infection to induce expression of ETS-1, a highly-conserved transcription factor (TF) that promotes restriction of A51R-deficient VV. Finally, we show that FACT also restricts unrelated cytoplasmic RNA viruses and that VV antagonism of FACT contributes to viral virulence in mice, establishing FACT as a critical component of antiviral immunity.

## Results

### VV A51R Interacts with the hSpt16 Subunit of Human FACT

To identify VV A51R-host interactions, we conducted a yeast two-hybrid screen with A51R bait and a human prey library. This screen identified hSpt16 as a top hit, with nine overlapping prey clones narrowing the putative A51R interaction domain to hSpt16 residues 758-893 within the middle domain (**Fig. 1a**). Co-immunoprecipitation (Co-IP) studies confirmed interaction between hSpt16 and Flag-A51R (FA51R) in A549 cells after infection with a VV strain expressing FA51R under its natural promoter (ΔA51R^FA51R^)^8^ (**Fig. 1bc**). Notably, SSRP1 did not Co-IP with A51R (**Fig. 1b**), indicating that A51R exclusively interacts with the hSpt16 subunit of FACT.

**Figure 1.**
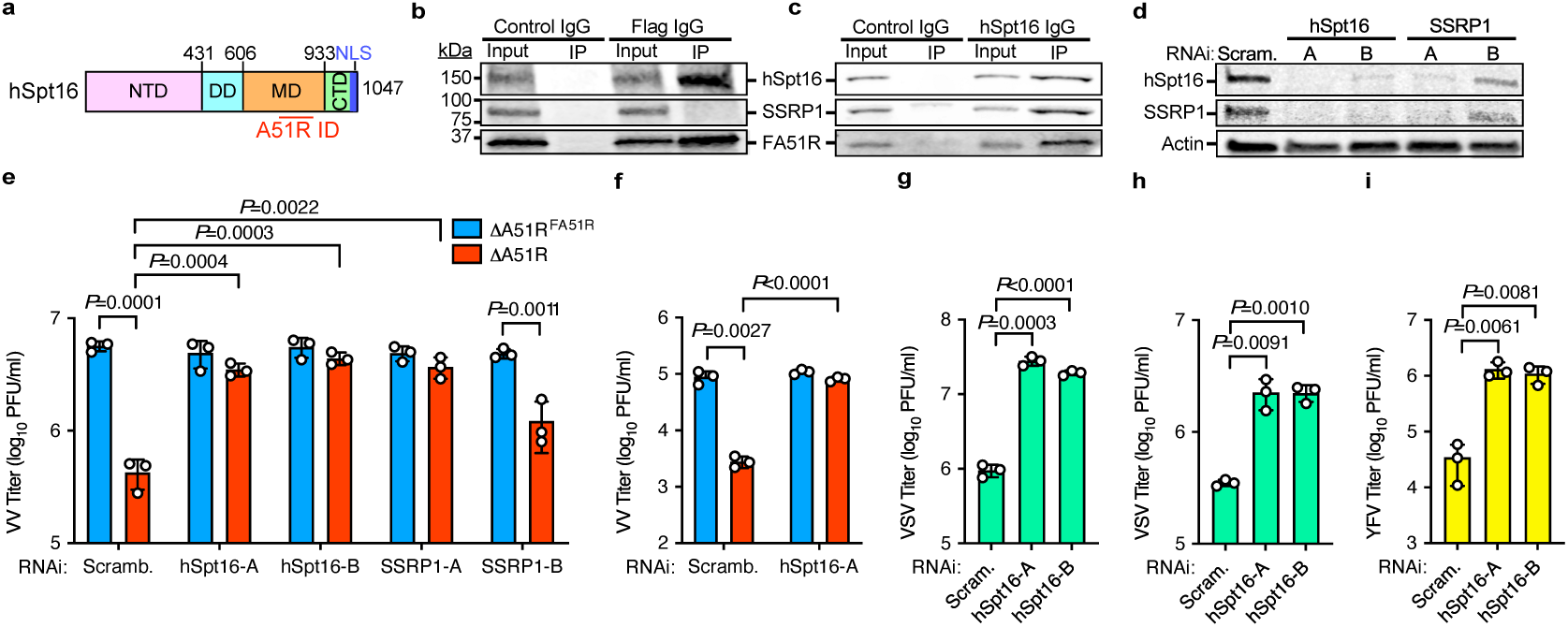
VV A51R interacts with hSpt16 and FACT depletion promotes cytoplasmic virus replication. **a**, Putative A51R interaction domain (ID) from yeast two-hybrid mapped onto hSpt16. NTD, N-terminal Domain; DD, Dimerization Domain; MD, Middle Domain; CTD, C-terminal Domain^21^. **b**,**c**, Immunoblots (IB) of reciprocal Co-IPs of endogenous hSpt16 with Flag-A51R (FA51R) in ΔA51R^FA51R^-infected A549 whole cell extract (WCE) using 10% protein gels. **d**, IB of A549 WCE 48 h after indicated RNAi. Scram., scrambled. **e**,**f**, VV titers 48 hpi [multiplicity of infection (MOI)=0.01)] in A549 (**e**) or NHDF (**f**) cell cultures transfected with indicated RNAi treatments as in (**d**). **g**,**h**, VSV-GFP^8^ titers 24 hpi (MOI=0.001) in A549 (**g**) and NHDF (**h**) cells transfected with indicated RNAi treatments as in (**d**). **i**, YFV-17D-Venus^38^ titers 24 hpi (MOI=0.01) in A549 cells transfected with indicated RNAi treatments as in (**d**). Data in **e-g** are means ± SD; n=3 and statistical significance was determined by unpaired two-tailed Student’s t-test. Only statistical comparisons with *P*<0.05 are shown.

### FACT Inhibits Cytoplasmic DNA and RNA Virus Replication

Since FACT had not been shown to regulate cytoplasmic virus replication, we next used RNA interference (RNAi) to deplete FACT in cells prior to infection with either ΔA51R^FA51R^ or A51R knockout (ΔA51R)^8^ strains to determine if FACT influences VV replication. It is important to note that under control RNAi treatments, the ΔA51R strain replicates to ~10-30-fold lower titers than the revertant ΔA51R^FA51R^ virus, consistent with the known replication defect of ΔA51R^8^. Also, prior work showed that mRNAs encoding hSpt16 and SSRP1 bind to the FACT subunit proteins themselves and stabilize the complex, thus RNAi targeting either hSpt16 or SSRP1 depletes both proteins^16^ (**Fig. 1d**). While FACT depletion did not affect ΔA51R^FA51R^ replication, multiple, independent hSpt16 or SSRP1 RNAi treatments enhanced ΔA51R replication to titers indistinguishable from ΔA51R^FA51R^ infections (**Fig. 1e**). Similar results were observed in primary neonatal human dermal fibroblast (NHDF) cells (**Fig. 1f**). These data suggest that FACT depletion complements the replication defect of A51R-deficient VV.

We also examined negative-sense [vesicular stomatitis virus (VSV)]^8^ and positive-sense [yellow fever virus (YFV)^17^] ssRNA viruses for changes in replication after FACT depletion. hSpt16 RNAi enhanced both VSV (**Fig. 1gh**) and YFV (**Fig. 1i**) replication, indicating that FACT broadly restricts cytoplasmic DNA and RNA viruses. We next focused on examining VV A51R-hSpt16 interactions in more detail to probe the role of FACT in antiviral immunity.

### Poxvirus A51R Proteins Specifically, and Directly, Bind the Middle Domain of a novel, SUMOylated form of hSpt16 Using a Conserved Motif

Prior studies have presented hSpt16 as a single band on immunoblots^16, 18^. Indeed, hSpt16 resolves as one band on 10% acrylamide gels (**Fig. 1bc**), but we observed two hSpt16 bands of ~140 and 155 kDa on 6% gels (**Fig. 2a**). Strikingly, A51R specifically bound only the larger hSpt16 form (**Fig. 2a**). The ~15-20 kDa difference between bands suggested that the upper band may be a novel singly-SUMOylated form of hSpt16. Thus, we re-probed our Co-IP membranes in a separate channel with SUMO-1 antibodies (Ab) and found them to specifically bind the upper hSpt16 band (**Fig. 2a**). Reciprocal Co-IPs with hSpt16 Ab confirmed enrichment of both hSpt16 forms and A51R interaction (**Fig. 2b**). To confirm hSpt16 SUMOylation, we treated cells with tannic acid (TA), a global SUMOylation inhibitor^19^. TA specifically depleted the upper hSpt16 band (**Fig. 2c**). In addition, only this upper hSpt16 band was enriched in SUMOylated protein fractions immunoprecipitated from cell extracts (**Fig. 2de**). Given the conservation of Spt16 among eukaryotes (**Extended Data Fig. 1a**), we assessed Spt16 SUMOylation in different human cell types and eukaryotic species. We found SUMOylated Spt16 in multiple human cell lines and primary NHDFs and in every monkey, mouse, hamster, rabbit, and bat cell line tested (**Extended Data Fig. 1bc**). Spt16 was also SUMOylated in insect cells and in *Caenorhabditis elegans* tissue extracts (**Extended Data Fig. 1d**), suggesting that this modification is conserved in invertebrates. From here, we refer to SUMOylated hSpt16 as “hSpt16^SUMO^”.

**Figure 2.**
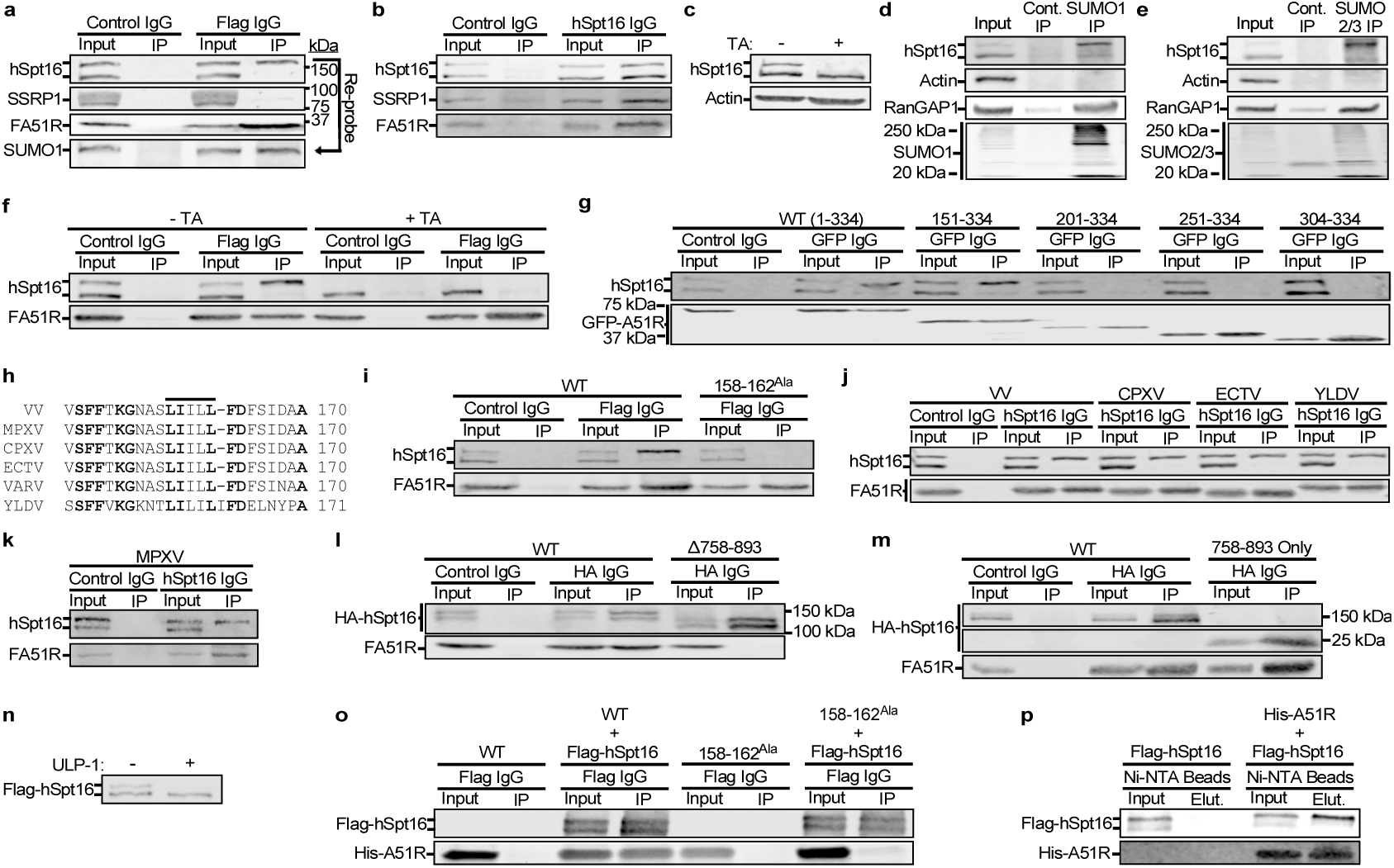
Poxvirus A51R proteins specifically, and directly, bind the middle domain of a novel, SUMOylated form of hSpt16 using a conserved motif. **a,b,** Reciprocal Co-IPs of hSpt16 with FA51R in ΔA51R^FA51R^-infected A549 WCE using 6% gels. **c,** hSpt16 IB of WCE from A549 cells treated with TA. **d,e,** IB of immunoprecipitated SUMO-1- (d) or SUMO-2/3- (e) conjugated protein fractions in A549 WCE. RanGAP1 is a known SUMOylated protein and used as a control^39^. **f,** hSpt16-FA51R Co-IP in ΔA51R^FA51R^-infected A549 WCE ± TA treatment. **g,** Co-IP of transfected VV GFP-A51R truncation mutants encoding indicated A51R residues with hSpt16 in 293T WCE. **h,** Conservation of hydrophobic motif (horizontal line) in poxvirus A51R proteins. **i,** Co-IP of transfected VV FA51R constructs with hSpt16 in 293T WCE. **j,** Co-IP of transfected Flag-tagged poxvirus A51R proteins with hSpt16 in 293T WCE. **k,** Co-IP of transfected VV FA51R and HA-hSpt16 constructs in 293T WCE. **l,** IB of HA-hSpt16-transfected 293T WCE ± TA treatment. **m,** Co-IP of transfected VV FA51R and HA-hSpt16 constructs in 293T WCE. **n,** IB of purified Flag-hSpt16 protein ± ULP-1 treatment. **o,** *in vitro* Co-IP of Flag-hSpt16 with WT or mutant His-A51R. **p,** *in vitro* nickel bead pulldown of WT His-A51R and Flag-hSpt16. Images in a-p are representative of at least two independent experiments.

To determine if hSpt16 SUMOylation is required for A51R interaction, we performed Co-IPs from VV-infected cells treated with TA. In the absence of TA, A51R pulled down hSpt16^SUMO^. However, interaction was lost after TA treatment (**Fig. 2f**), indicating that hSpt16 SUMOylation is required for A51R binding. Next, we used GFP-tagged VV A51R truncation mutants to identify a region between A51R a.a. 151-201 required for hSpt16^SUMO^ interaction (**Fig. 2g**). Alignment of this region with other poxvirus A51R proteins sharing 35-95% a.a. identity to VV A51R (**Extended Data Fig. 1e**) revealed a conserved hydrophobic motif (VV A51R a.a. 158-162) (**Fig. 2h**). Substitution of motif residues with alanine (A51R^158-162Ala^) prevented VV A51R-hSpt16^SUMO^ interaction (**Fig. 2i**), indicating a role for this motif in this interaction. Given the conservation of this motif, we tested ectromelia virus (ECTV), cowpox virus (CPXV), Yaba-like disease virus (YLDV), and Mpox virus (MPXV) A51R proteins for interaction with hSpt16^SUMO^ and found all four to bind hSpt16^SUMO^ (**Fig. 2jk**). Thus, A51R-Spt16^SUMO^ binding is a conserved poxvirus-host interaction.

We next determined if the putative A51R interaction domain in hSpt16 identified by yeast two-hybrid (a.a. 758-893; **Fig. 1a**) was required for A51R-hSpt16^SUMO^ Co-IP. Deletion of this domain from HA-tagged hSpt16 prevented Co-IP with A51R (**Fig. 2l**), confirming its role in A51R interaction. Surprisingly, this deletion mutant still ran as two bands on immunoblots suggesting it is still SUMOylated (**Fig. 2l**). These data suggested that A51R may not interact with hSpt16 through the SUMO moiety itself but rather SUMOylation may cause a conformational change in hSpt16 that exposes the A51R binding site. Consistent with this, a fragment encoding only hSpt16 residues 758-893 ran as a single band on immunoblots but still interacted with A51R (**Fig. 2m**).

We next asked if purified, His-tagged A51R (His-A51R) protein directly interacts with Flag-tagged hSpt16 (Flag-hSpt16) purified from human cells. We confirmed that our Flag-hSpt16 protein was SUMOylated with ULP-1 protease treatment, which cleaves SUMO moieties^20^ (**Fig. 2n**). Using reciprocal *in vitro* Co-IPs and pulldowns, we found His-A51R to Co-IP with Flag-hSpt16 in Flag Ab IPs and only the SUMOylated form of Flag-hSpt16 to bind His-A51R in nickel bead pulldowns (**Fig. 2op**), indicating a direct interaction. Notably, a His-A51R^158-162Ala^ mutant did not Co-IP with Flag-hSpt16 (**Fig. 2o**).

### VV A51R Tethers hSpt16^SUMO^ to MTs to Block hSpt16^SUMO^ Nuclear Accumulation Triggered by Early VV Gene Expression

In cells, VV A51R exclusively localizes to cytosolic MTs^8^ while hSpt16 functions in the nucleus^21^. Thus, we asked if A51R alters hSpt16 localization. Using a cell line expressing GFP-tagged hSpt16 (GFP-hSpt16), we found a portion of GFP-hSpt16 to colocalize with A51R in the cytosol in ΔA51R^FA51R^-infected cells. In contrast, no such cytosolic GFP-hSpt16 enrichment was found in mock- or ΔA51R-infected cells (**Fig. 3a**). This suggested that A51R may tether hSpt16^SUMO^ to MTs. In another study (Seo et al., in prep), we identified a C-terminal A51R domain (a.a. 254-302) that resembles the MT-binding domain of Tau, a cellular MT-associated protein^22^ (**Extended Data Fig. 2a**). A “triple” A51R mutant encoding R275A/K295A/K302A substitutions in this domain cannot colocalize with, or bundle, MTs or protect MTs from depolymerization by nocodazole, unlike WT A51R (**Extended Data Fig. 2b**). The A51R^Triple^ mutant is also unable to co-sediment with MTs *in vitro* (**Fig. 3bc**), indicating these substitutions destroy direct A51R-tubulin interactions. However, A51R^158-162Ala^ still co-sediments with MTs *in vitro* (**Fig. 3d**) and A51R^Triple^ still interacts with hSpt16^SUMO^ (**Fig. 3e**). Moreover, hSpt16 only interacted with tubulin when in the presence of A51R *in vitro* (**Fig. 3fg**). These data suggest that A51R tethers hSpt16^SUMO^ to MTs by simultaneously binding hSpt16^SUMO^ and tubulin through distinct domains.

**Figure 3.**
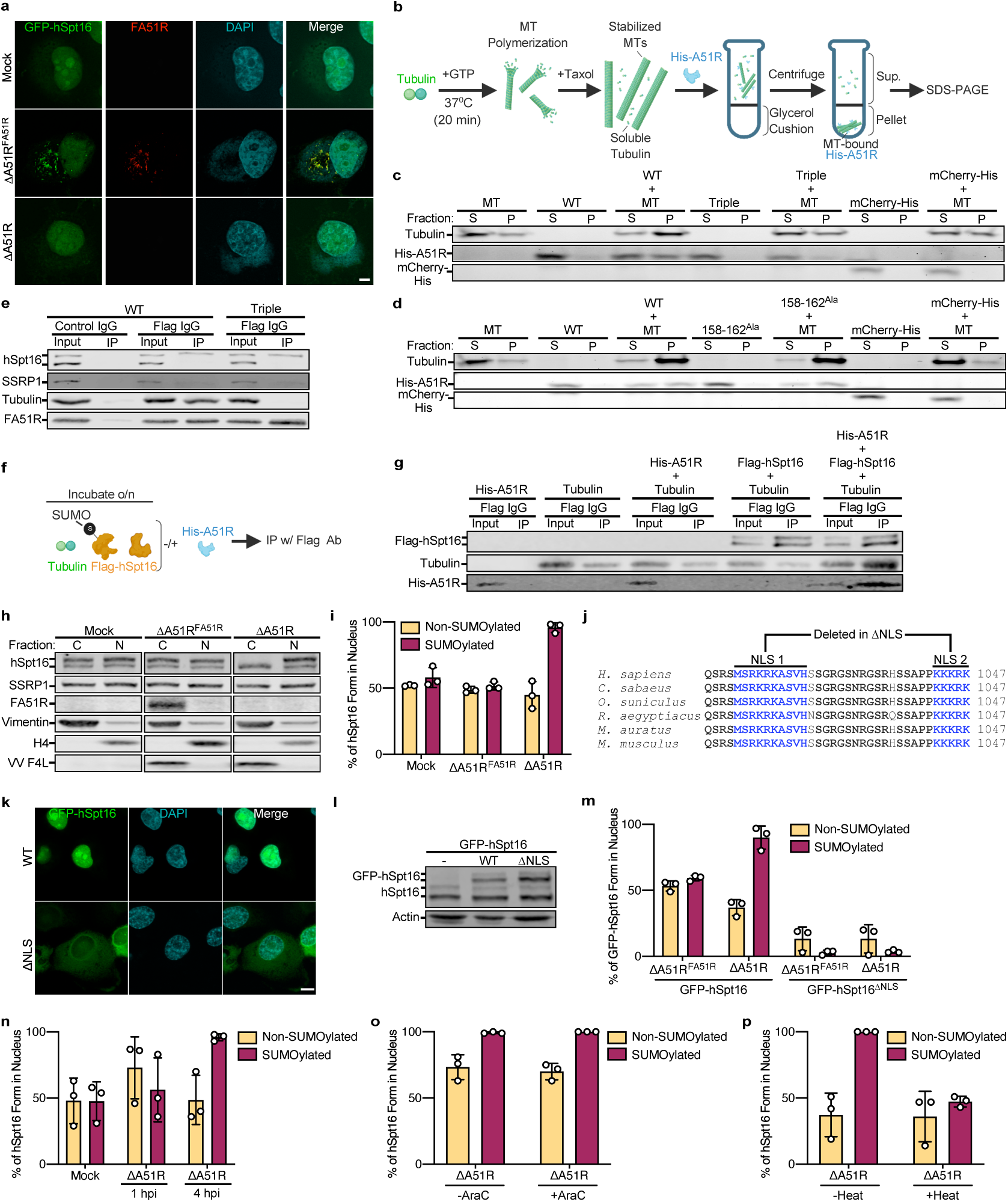
VV A51R tethers hSpt16^SUMO^ to MTs to block hSpt16^SUMO^ nuclear accumulation triggered by VV early gene expression. **a**, Immunofluorescence (IF) images of GFP-hSpt16-expressing U2OS cells 18 hpi with VV strains. Cytosolic DAPI staining marks VV infection. Scale bar, 5 μm. **b**, Overview of *in vitro* MT co-sedimentation assay. **c**,**d**, *in vitro* MT co-sedimentation assays with purified WT and mutant His-A51R. **e**, Co-IP of transfected WT and mutant FA51R constructs with hSpt16 and tubulin in 293T WCE. **f**, Overview of *in vitro* Flag-hSpt16 Co-IP with tubulin in the absence/presence of His-A51R. **g**, *in vitro* Co-IP of purified Flag-hSpt16, His-A51R and tubulin. **h**,**i**, Representative IB (**h**) and quantification (**i**) of U2OS cytoplasmic (C) and nuclear (N) fractions 18 hpi with VV strains. VV F4L marks infection. Data in **i** are means ± SD; n=3. **j**, hSpt16 NLS motifs deleted in ΔNLS mutant and their conservation in eukaryotic Spt16 proteins. **k**, Representative IF images of WT or ΔNLS GFP-hSpt16 U2OS cells. Scale bar, 10 μm. **l**, IB of WCE from U2OS cells expressing WT or ΔNLS GFP-hSpt16 proteins. **m**, Quantification of fractionation experiments of WT or ΔNLS GFP-hSpt16-expressing U2OS cells infected with VV strains for 18 h. Data are means ± SD; n=3. **n**, Quantification of fractionation experiments of A549 cells mock- or ΔA51R-infected for indicated times. Data are means ± SD; n=3. **o**, Quantification of fractionation experiments of U2OS cells infected with ΔA51R strain for 4 h. Where indicated, AraC was added to media 1 hpi. Data are means ± SD; n=3. **p,** Quantification of fractionation experiments of U2OS cells infected with ΔA51R strain for 4 h. Where indicated, ΔA51R was heat-inactivated prior to infection. Data are means ± SD; n=3. Images in **a, c-e, g-h,** and **k-l** are representative of at least two independent experiments.

We next used cell fractionation to ask if the intracellular distribution of hSpt16 is altered during VV infection. Strikingly, hSpt16^SUMO^ was specifically absent from cytosolic fractions in ΔA51R-infected cells and only present in nuclear fractions, in contrast to mock- and ΔA51R^FA51R^-infected cells (**Fig. 3hi**). This suggested that ΔA51R infection either triggers changes in hSpt16 SUMOylation, hSpt16^SUMO^ stability, or hSpt16^SUMO^ cytosolic/nuclear distribution and that A51R blocks these infection-induced changes. However, total hSpt16^SUMO^ levels did not overtly change in ΔA51R-infected cells over a 24 h time-course (**Extended Data Fig. 3**). To examine changes in hSpt16 nuclear localization, we first identified the hSpt16 nuclear localization signal (NLS) using NLS prediction software^23, 24^. We identified two conserved NLS motifs near the C-terminus of hSpt16 (**Fig. 3j**) that, when deleted, prevented GFP-hSpt16 nuclear import (**Fig. 3k**). We then generated a GFP-hSpt16^ΔNLS^-expressing cell line and found GFP-hSpt16^ΔNLS^ to still be SUMOylated, indicating that SUMOylation occurs in the cytosol, but it remained in the cytosol after ΔA51R infection (**Fig. 3lm**). These data suggest that A51R inhibits infection-triggered hSpt16^SUMO^ nuclear accumulation.

To determine which step of the VV life cycle was required for hSpt16^SUMO^ nuclear accumulation, we first analyzed the timing of hSpt16^SUMO^ cytosolic/nuclear distribution changes after ΔA51R infection. We found hSpt16^SUMO^ nuclear accumulation to occur by 4 hpi (**Fig. 3n**), a time at which VV is known to initiate VV DNA replication and late gene expression. Since VV DNA replication occurs prior to late gene expression, we next analyzed hSpt16^SUMO^ cytosolic/nuclear distribution during ΔA51R infection in the presence of cytosine arabinoside (AraC), a VV DNA replication inhibitor^8^. Interestingly, AraC did not block ΔA51R-induced nuclear accumulation of hSpt16^SUMO^ (**Fig. 3o**), suggesting that a step in the VV life cycle prior to VV DNA replication such as entry or early gene expression triggers hSpt16^SUMO^ nuclear accumulation. Thus, we employed a heat inactivation protocol that allows virion entry, but inactivates early VV gene expression^8^. Heat treatment completely abrogated ΔA51R-induced hSpt16^SUMO^ nuclear accumulation, suggesting that early VV gene expression is required to trigger this host response (**Fig. 3p**).

### VV A51R Outcompetes SSRP1 to Inhibit hSpt16^SUMO^-SSRP1 Interaction

To ask if A51R affects hSpt16-SSRP1 interaction, we examined GFP-hSpt16-SSRP1 Co-IP in U2OS cells expressing GFP-hSpt16 during mock, ΔA51R^FA51R^, or ΔA51R infection. Compared to mock- and ΔA51R-infected lysates, the amount of SSRP1 bound to GFP-hSpt16 was reduced in ΔA51R^FA51R^ lysates (**Fig. 4a**). SSRP1 Ab Co-IPs in parental U2OS cells also showed reduced hSpt16^SUMO^-SSRP1 interaction in ΔA51R^FA51R^ lysates (**Fig. 4b**). This implied that A51R competes with SSRP1 for hSpt16^SUMO^ binding, so we examined the ability of His-A51R and His-tagged SSRP1 (His-SSRP1) to compete for Flag-hSpt16 binding *in vitro*. We first confirmed that purified Flag-hSpt16 and His-SSRP1 proteins interacted as expected using reciprocal pulldowns (**Fig. 4cd**). Of note, we did not observe differences in His-SSRP1 affinity for SUMOylated/SUMOless Flag-hSpt16 (**Fig. 4d**). We next incubated increasing amounts of His-SSRP1 with preformed His-A51R-Flag-hSpt16^SUMO^ complexes and then conducted SSRP1 Ab Co-IPs to assess the relative amounts of Flag-hSpt16^SUMO^ bound to His-SSRP1. Even at a 10-fold molar excess of His-SSRP1:His-A51R, Flag-hSpt16^SUMO^ did not interact with His-SSRP1 in the presence of His-A51R. However, His-SSRP1 interacted efficiently with Flag-hSpt16^SUMO^ in the absence of His-A51R (**Fig. 4ef**), suggesting that His-SSRP1 poorly competes with His-A51R for Flag-hSpt16^SUMO^. Consistent with this, when increasing amounts of His-A51R were added to preformed His-SSRP1-Flag-hSpt16 complexes, His-SSRP1-Flag-hSpt16 interaction decreased with increasing His-A51R (**Fig. 4gh**). These data suggest that A51R can outcompete with SSRP1 for binding to hSpt16^SUMO^ to disrupt FACT complex formation.

**Figure 4.**
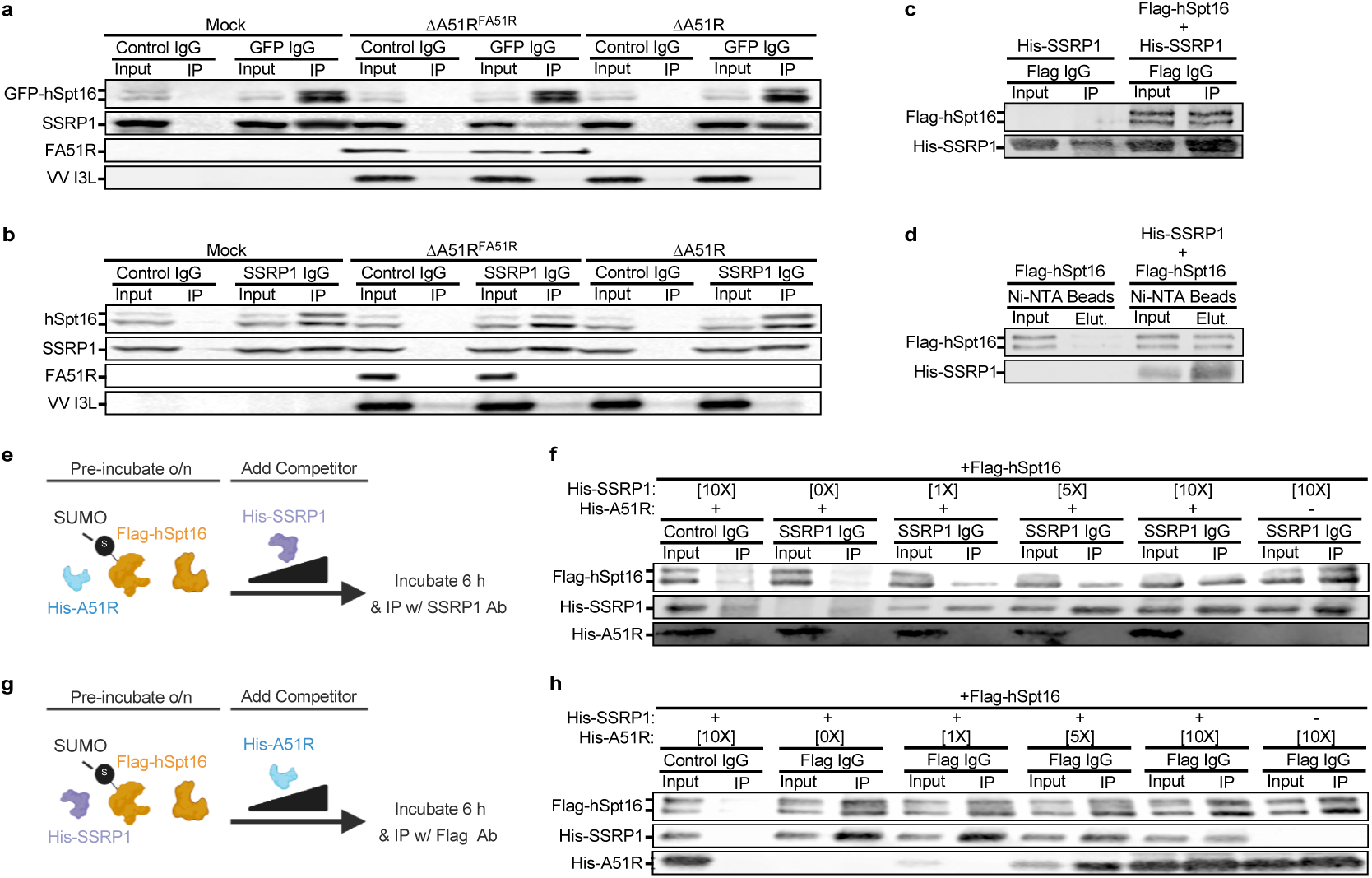
VV A51R outcompetes SSRP1 to inhibit hSpt16^SUMO^-SSRP1 interaction. **a**, IB of GFP-hSpt16-SSRP1 Co-IP in GFP-hSpt16-expressing U2OS WCE 18 hpi with VV strains. VV I3L marks infection. **b**, IB of hSpt16-SSRP1 Co-IP in U2OS WCE 18 hpi with VV strains. **c**,**d**, *in vitro* Co-IP (**c**) and pulldown (**d**) assays with purified Flag-hSpt16 and His-SSRP1. **e**-**h**, *in vitro* competition assays with preformed complexes of Flag-hSpt16-His-A51R (**e,f**) or Flag-hSpt16-His-SSRP1 (**g,h**) incubated with increasing molar ratios of His-SSRP1 (**e,f**) or His-A51R (**g,h**) and then subjected to SSRP1 Ab IP (**e,f**) or Flag Ab IP (**g,h**). Images in **a**-**f** are representative of at least two independent experiments. Images in **e** and **g** were made with biorender.com.

### hSpt16^SUMO^ Nuclear Localization is Required for Virus Restriction

To determine if hSpt16^SUMO^ nuclear localization is required for VV restriction, we infected GFP-hSpt16- or GFP-hSpt16^ΔNLS^-expressing cells with ΔA51R^FA51R^ or ΔA51R strains after RNAi-mediated depletion of endogenous hSpt16. Normally, hSpt16 RNAi depletes both hSpt16 and SSRP1 because of the stabilization of FACT subunits by their own mRNAs. However, expression of GFP-hSpt16 mRNA can stabilize SSRP1 protein levels^16^. Thus, we used hSpt16 RNAi designed to target the endogenous *hSpt16* gene (but not the *GFP-hSpt16* gene) to specifically deplete endogenous hSpt16 proteins while retaining SSRP1 levels (**Fig. 5a**). Interestingly, ΔA51R replicated to higher titers in cells expressing GFP-hSpt16^ΔNLS^, but not WT GFP-hSpt16, after endogenous hSpt16 depletion (**Fig. 5b**), suggesting that hSpt16^SUMO^ nuclear import is required for its antiviral function.

**Figure 5.**
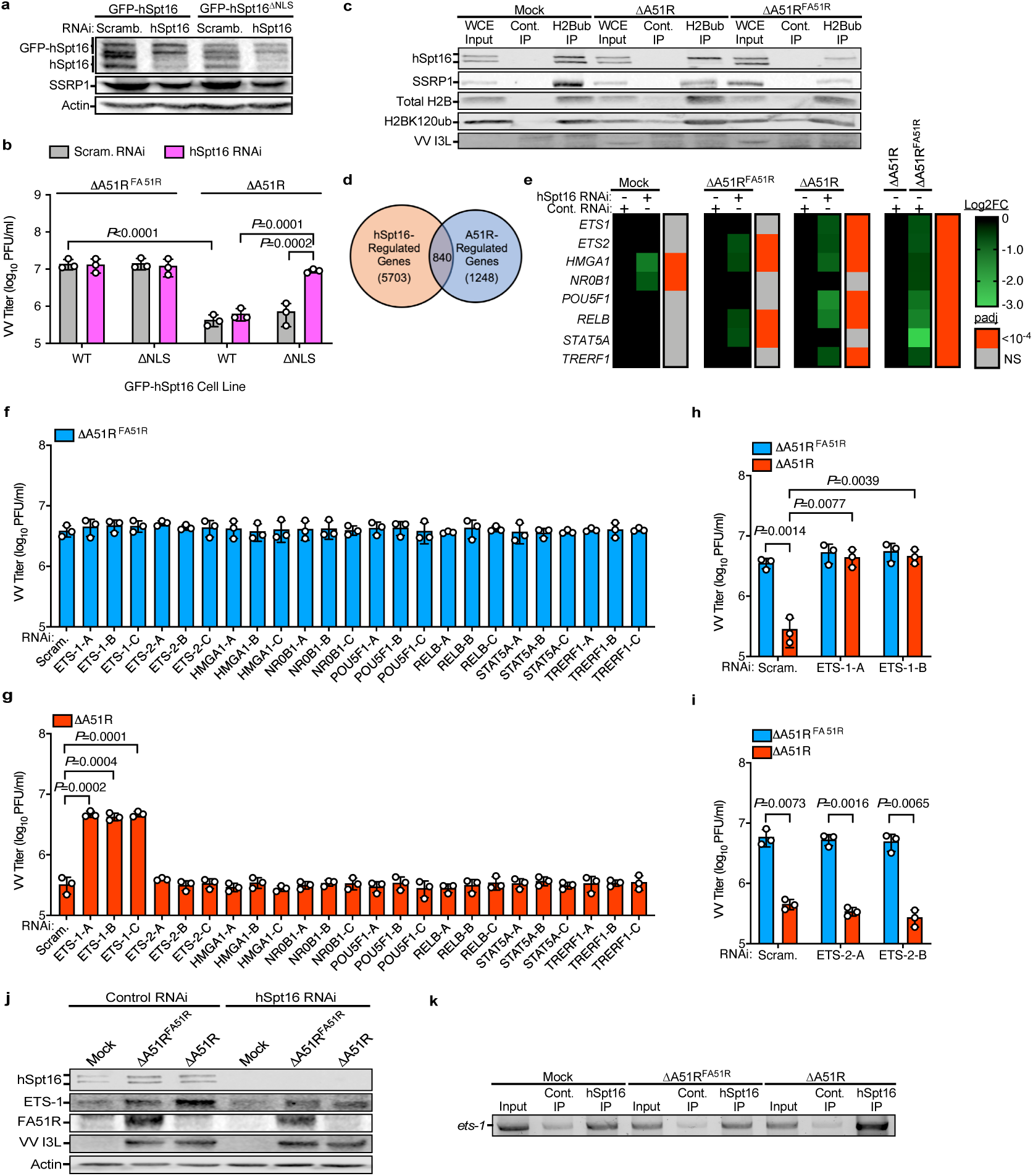
hSpt16^SUMO^-SSRP1 (FACT) complexes bind transcriptionally active chromatin during infection to activate ETS-1 expression and VV restriction, which is antagonized by VV A51R. **a,** IB of U2OS WCE expressing WT or ΔNLS GFP-hSpt16 48 h post-RNAi. **b,** VV titers 48 hpi (MOI=0.01) of cell lines treated as in (b). Data are means ± SD; n=3. **c,** IB of Co-IP of FACT subunits with H2BK120ub in U2OS nuclear extracts 18 hpi with VV strains. WCE inputs indicate total protein levels prior to nuclear protein extraction and equal division into control or H2BK120ub Ab IPs. **d,** Non-proportional Venn diagram showing total human DEGs after hSpt16 RNAi across mock- and VV-infected samples and overlap with DEGs between ΔA51R^FA51R^ and ΔA51R infections in control RNAi conditions. **e,** Heat map of relative RNA-seq expression level of immunity-related TFs found to be FACT/A51R-regulated genes under indicated RNAi/infection conditions. Heat maps indicate relative log2-fold change (decrease) and adjusted *P* values. **f,g,** VV titers 48 hpi for TF RNAi screens using ΔA51R^FA51R^ (f) and ΔA51R (g) in A549 cells (MOI=0.01). Data are means ± SD; n=3. **h,i,** VV titers in A549 cells 48 hpi (MOI=0.01) after ETS-1 (h) or ETS-2 (i) RNAi. Data are means ± SD; n=3. **j,** IB of ETS-1 in WCE of control or hSpt16-shRNA-expressing A549 cell lines 4 hpi with VV strains (MOI=10). **k,** Agarose gel image of ChIP-PCR assay of *ets-1* promoter-proximal region^31^ after control or hSpt16 Ab-based ChIP in A549 cells under infection conditions as in (j). Images in a, c, and j-k, are representative of at least two independent experiments. Statistical significance in f-i was determined by unpaired two-tailed Student’s t-test. Only statistical comparisons with *P*<0.05 are shown.

### Only SUMOylated hSpt16 Binds Transcriptionally Active Chromatin During Infection

Since FACT interacts with nucleosomes to regulate gene expression, we asked if hSpt16 SUMOylation influenced such interactions. However, no differences were found in the ability of Flag-hSpt16/hSpt16^SUMO^ to interact with purified histone complexes *in vitro* (**Extended Data Fig. 4a**). Furthermore, deSUMOylation of purified FACT with ULP-1 did not alter FACT binding to reconstituted (H3-H4)_2_ tetrasomes or (H2A-H2B)-(H3-H4)_2_ hexasomes *in vitro* (**Extended Data Fig. 4b**). Prior work showed FACT to associate with K120-monoubiquitinated H2B (H2BK120ub)^25, 26^, which marks active transcription sites in chromatin^15^. Since H2B proteins used in our *in vitro* assays were purified from bacteria, they lack ubiquitination. Therefore, we examined hSpt16/hSpt16^SUMO^ binding to H2BK120ub in nuclear extracts. In the absence of VV infection, both hSpt16 forms bound H2BK120ub. However, only hSpt16^SUMO^ bound H2BK120ub during infection and this interaction was reduced during ΔA51R^FA51R^ infection (**Fig. 5c**). This suggests that hSpt16 requires SUMOylation to bind H2BK120ub during infection and A51R inhibits this interaction.

### VV A51R Inhibits FACT-Dependent Expression of an Antiviral Transcription Factor

The previous results led us to hypothesize that hSpt16^SUMO^ activates FACT-dependent antiviral gene expression to restrict virus replication. To identify cellular genes regulated by FACT during infection, we extracted total RNA from A549 cells expressing control or hSpt16-targeted short hairpin RNA after 4 h of mock, ΔA51R^FA51R^, or ΔA51R infection, and used mRNA-sequencing (RNA-seq) to identify differentially expressed genes (DEGs) after FACT depletion. We chose 4 hpi as our time point because hSpt16^SUMO^ nuclear accumulation occurs by that time (**Fig. 3n**).

Between mock-, ΔA51R^FA51R^- and ΔA51R-infected treatments, we identified 5703 DEGs after hSpt16 RNAi in at least one of these three infection conditions (**Figure 5d**; **Supplementary Table 1a-e**). We then cross-referenced these “hSpt16-regulated genes” with the 1248 DEGs between ΔA51R^FA51R^ and ΔA51R infections in control RNAi cells (“A51R-regulated genes”) (**Supplementary Table 1f**). This identified 840 genes that are differentially expressed after hSpt16 RNAi and in the presence of A51R expression (**Fig. 5d**; **Supplementary Table 1g**). We reasoned that A51R likely inhibits FACT-dependent expression of immunity genes, so we analyzed these 840 genes for immunity-related gene ontology classifications and identified 76 known immunity genes that were down-regulated in both hSpt16 RNAi treatments and in the presence of A51R. Notably, only ~6% of these 76 genes are IFN-stimulated genes in A549 cells^27, 28^ (**Supplementary Table 1h**), suggesting that FACT-induced antiviral responses are distinct from the IFN pathway.

As master regulators of transcription, we suspected that TFs were likely involved in FACT-dependent antiviral responses. Among the identified 76 hSpt16/A51R-regulated immunity genes were 8 TFs (**Fig. 5e**). Using RNAi, we screened these TFs for potential antiviral function during ΔA51R^FA51R^ or ΔA51R infection and identified ETS-1 as our only hit. ETS-1 is a member of the evolutionarily-conserved “E26 transformation specific (ETS)” TF family^29^ and was recently implicated in the immune response to bacterial infection^30^, but it was unclear if ETS-1 also functions in antiviral immunity. Interestingly, ETS-1 RNAi enhanced ΔA51R, but not ΔA51R^FA51R^, replication (**Fig. 5fg**). In side-by-side RNAi experiments with both VV strains, ETS-1 RNAi enhanced ΔA51R replication to levels that were indistinguishable from ΔA51R^FA51R^ (**Fig. 5h**), suggesting that like hSpt16 RNAi (**Fig. 1ef**), ETS-1 knockdown complements ΔA51R replication defects. Notably, the related ETS-2 TF was also hSpt16/A51R-regulated (**Fig. 5e**), but its knockdown did not affect VV replication (**Fig. 5fg,i**).

To determine the VV life cycle step required to trigger ETS-1 expression, we first analyzed the timing of ETS-1 expression after VV infection. We found ETS-1 induction during ΔA51R infection to occur by 4 hpi (**Extended Data Fig. 5a**) and to require early VV gene expression (**Extended Data Fig. 5b**), but not VV DNA replication (**Extended Data Fig. 5c**), which is consistent with the timing and requirements for ΔA51R-triggered hSpt16^SUMO^ nuclear accumulation (**Fig. 3n-p**). We next determined if ETS-1 protein levels change during VV infection in the absence or presence of hSpt16 RNAi. Consistent with our RNA-seq, ETS-1 was induced during virus infection in an hSpt16-dependent manner. However, stronger ETS-1 induction occurred in ΔA51R infections (**Fig. 5j**), suggesting this mutant has a reduced ability to block ETS-1 expression. Notably, no differences in IFN pathway activation were noted when comparing ΔA51R versus ΔA51R^FA51R^ infections under control or hSpt16 RNAi conditions (**Extended Data Fig. 6**), supporting the idea that FACT-induced antiviral responses are distinct from the IFN pathway.

**Figure 6.**
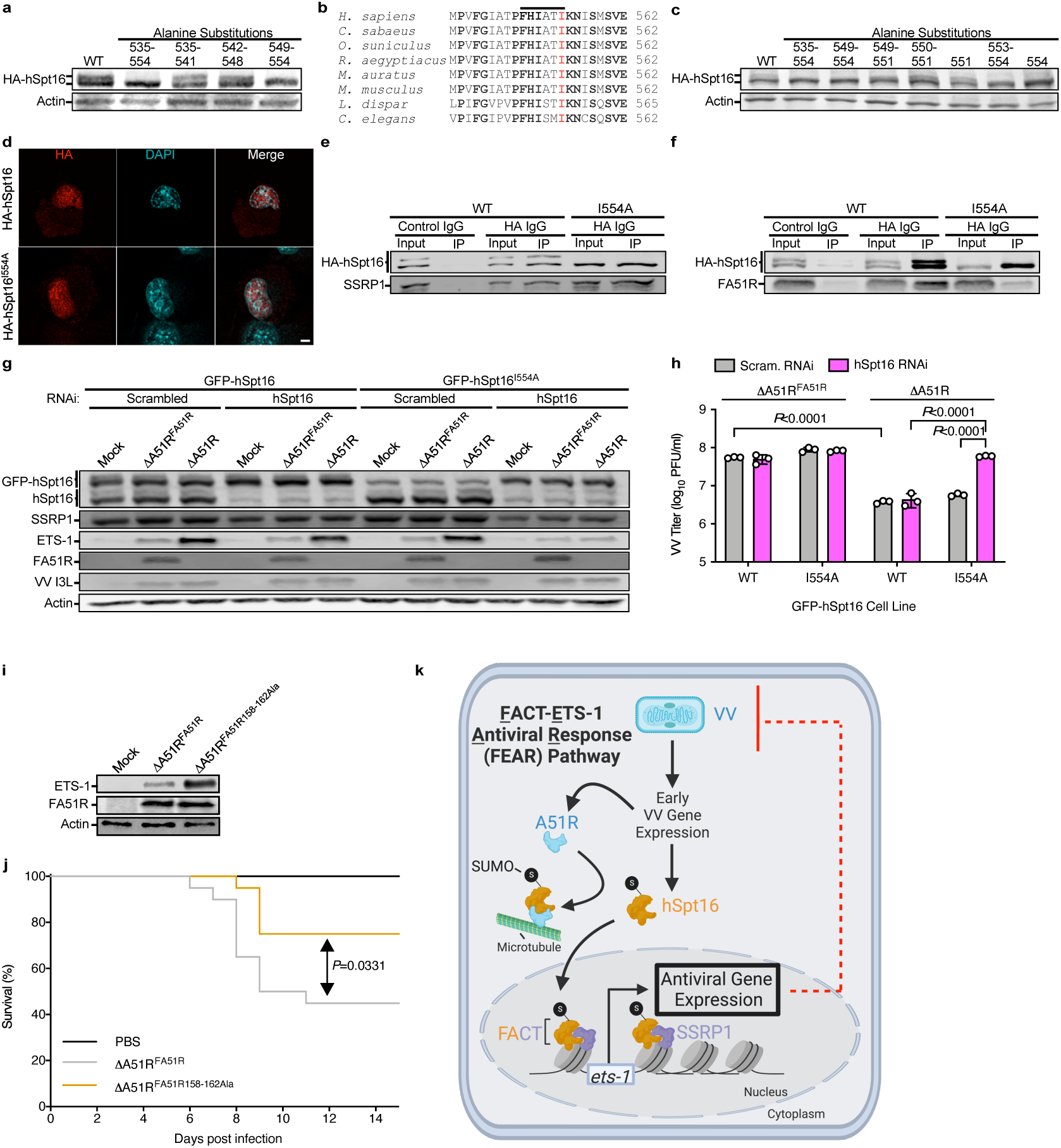
hSpt16^SUMO^ is required for ETS-1 expression and virus restriction, VV A51R-hSpt16^SUMO^ interaction suppresses ETS-1 expression and promotes VV virulence, and model for <u>F</u>ACT-<u>E</u>TS-1 <u>A</u>ntiviral <u>R</u>esponse (FEAR) pathway. a, IB of 293T WCE transfected with WT or mutant HA-hSpt16 constructs encoding alanine substitutions throughout indicated hSpt16 a.a. regions. b, Conservation of hSpt16 motif required for SUMOylation (horizontal line). c, IB of 293T WCE transfected with WT or mutant HA-hSpt16 constructs encoding alanine substitutions at indicated hSpt16 a.a. from (b). d, IF images of transfected WT and I554A mutant HA-hSpt16 constructs in U2OS cells. e, IB of Co-IP of transfected WT and mutant HA-hSpt16 constructs with SSRP1 in 293T WCE. f, IB of Co-IP of transfected WT and mutant HA-hSpt16 constructs with co-transfected FA51R constructs in 293T WCE. g, IB of U2OS WCE from cells stably expressing WT or I554A mutant GFP-hSpt16 that were transfected with scrambled or hSpt16 siRNAs for 48 h and then infected with indicated VV (MOI=10) for 4 h. h, VV titers 48 hpi (MOI=0.01) in cell lines from g after indicated RNAi. Data are means ± SD; n=3. i, IB of A549 WCE from cells infected with indicated VV (MOI=10) for 4 h. j, Percent mice (n=20 over two independent experiments) alive after VV infection. k, Model for <u>F</u>ACT-<u>E</u>TS-1 <u>A</u>ntiviral <u>R</u>esponse (FEAR) Pathway. VV early gene expression triggers hSpt16^SUMO^ nuclear accumulation and formation of FACT complexes containing hSpt16^SUMO^ that license FACT to associate with H2BK120ub sites in transcriptionally active chromatin and activate ETS-1 expression. ETS-1 subsequently activates antiviral gene expression to restrict VV replication. VV A51R is expressed early during infection^8^ and inhibits FEAR pathway by tethering hSpt16^SUMO^ to MTs and blocking its nuclear import. A51R may also prevent FACT complex formation in the cytosol by outcompeting SSRP1 for hSpt16^SUMO^ binding, which would also serve to prevent nuclear accumulation of hSpt16^SUMO^-containing FACT complexes (not shown). Image created with BioRender.com. Images in a-g, and i are representative of at least two independent experiments. Statistical significance in h was determined by unpaired two-tailed Student’s t-test and by Log-Rank (Mantel-Cox) tests in j. Only statistical comparisons with *P*<0.05 are shown.

A prior chromatin immunoprecipitation (ChIP)-sequencing study found hSpt16 and SSRP1 to bind the *ets-1* gene, with strongest binding peaks occurring in a promoter-proximal region^31^. Thus, we PCR-amplified a DNA fragment encompassing this *ets-1* promoter-proximal region after ChIP with hSpt16 Ab to determine if ETS-1 expression during infection correlated with hSpt16 binding to the *ets-1* gene. We found similar hSpt16 binding to *ets-1* in mock- and ΔA51R^FA51R^ infections but enhanced binding during ΔA51R infection (**Fig. 5k**), suggesting that FACT binds the *ets-1* locus during infection to promote expression and virus restriction while A51R antagonizes *ets-1* expression.

### hSpt16^SUMO^ is Required for ETS-1 Expression and Virus Restriction

We hypothesized that ETS-1 expression requires hSpt16^SUMO^ because A51R binds hSpt16^SUMO^ and inhibits ETS-1 induction. Thus, we sought to identify the hSpt16 SUMOylation site to construct a SUMOless mutant to test this hypothesis. However, after systematically converting all lysine (K) residues within ~50-200 a.a. stretches across HA-hSpt16 to alanine (A), no SUMOless mutants were found (**Extended Data Fig. 7ab**), but combining all 33 K to A substitutions across a.a. 457-830, produced a SUMOless hSpt16 protein (**Extended Data Fig. 7c**). Thus, hSpt16 appears to have several potential SUMOylation sites within a.a. 457-830 but these sites are likely mutually exclusive since we only ever observe mono-SUMOylated hSpt16 species on immunoblots.

Given the many K to A substitutions required to produce SUMOless hSpt16, we were concerned about pleiotropic effects on hSpt16 folding or function. Thus, we screened for alternative SUMOless mutants and identified a small motif (a.a. 549-554) within the hSpt16 dimerization domain required for SUMOylation (**Fig. 6a**). This motif does not contain K residues, but it is well-conserved among eukaryotes (**Fig. 6b**) and may interact with SUMOylation machinery. Notably, alanine substitutions of nearby K residues flanking this motif did not affect hSpt16 SUMOylation (**Extended Data Fig. 7d**). Further alanine mutagenesis within this motif revealed I554 to be required for hSpt16 SUMOylation (**Fig. 6c**). An HA-hSpt16^I554A^ mutant was still capable of nuclear localization (**Fig. 6d**) and SSRP1 interaction (**Fig. 6e**). However, it was severely impaired in A51R binding (**Fig. 6f**). Using GFP-hSpt16^I554A^-expressing cell lines, we next examined ETS-1 expression during mock-, ΔA51R^FA51R^-and ΔA51R-infections when endogenous hSpt16 was depleted by RNAi (as in **Fig. 5a**). In GFP-hSpt16-expressing cells, ΔA51R induced higher ETS-1 expression compared to ΔA51R^FA51R^ as expected, but this induction was largely abrogated in GFP-hSpt16^I554A^-expressing cells after knockdown of endogenous hSpt16 (**Fig. 6g**). Furthermore, loss of ETS-1 expression was concomitant with increases in ΔA51R replication to titers comparable to ΔA51R^FA51R^ (**Fig. 6h**). Collectively, these data suggest that hSpt16^SUMO^ is required for ETS-1 expression and virus restriction.

### VV A51R-hSpt16^SUMO^ Interaction Inhibits ETS-1 Expression and Promotes VV Virulence

Next, we asked if A51R-hSpt16^SUMO^ interaction contributes to ETS-1 expression inhibition by VV. Therefore, we constructed a VV revertant strain encoding the hSpt16^SUMO^ interaction-deficient A51R^158-162Ala^ mutant (ΔA51R^FA51R158-162Ala^) and compared its ability to suppress ETS-1 induction to ΔA51R^FA51R^. The ΔA51R^FA51R158-162Ala^ mutant displayed a reduced ability to antagonize ETS-1 expression, suggesting that A51R-hSpt16^SUMO^ interaction inhibits ETS-1 induction after infection (**Fig. 6i**). Finally, we asked if FACT antagonism influences VV pathogenesis in mice by comparing the virulence of ΔA51R^FA51R158-162Ala^ and ΔA51R^FA51R^ strains after intranasal inoculation. Mice inoculated with A51R^158-162Ala^ exhibited greater survival than ΔA51R^FA51R^-infected animals (**Fig. 6j**), indicating that FACT antagonism promotes VV virulence.

## Discussion

Our study reveals that early VV gene expression triggers FACT-dependent ETS-1 expression that subsequently restricts viral replication. We term this host response the “<u>F</u>ACT-<u>E</u>TS-1 <u>A</u>ntiviral <u>R</u>esponse (FEAR)” Pathway. Central to the activation of this pathway is hSpt16 SUMOylation, which licenses FACT to bind transcriptionally active chromatin during infection. Given that Spt16 SUMOylation was wide spread among the mammalian and invertebrate species tested, SUMOylation may be an ancient mechanism to regulate antiviral FACT function. The physiological relevance of the FEAR pathway is underscored by our finding that VV A51R counters this pathway by competing with SSRP1 to bind hSpt16^SUMO^ and tether it to MTs to block its nuclear accumulation (**Fig. 6k**). Moreover, the reduced pathogenicity of VV strains encoding A51R substitutions that prevent hSpt16^SUMO^ interaction, highlight the contribution of this virus-host interaction to viral disease. Given that A51R proteins from other poxviruses (e.g. ECTV, CPXV, YLDV, etc.) all bind hSpt16^SUMO^ (**Fig. 2jk**) and associate with MTs^8^, it is clear that FACT antagonism is an important poxvirus immune evasion mechanism.

Inducible transcriptional responses to infection have long been recognized as integral aspects of antiviral immunity^32–34^. However, prior studies have primarily focused on the role of TFs in activating such responses and on the IFN pathway in particular^35^. Our work suggests that both chromatin remodeling proteins (FACT) and TFs (ETS-1) serve critical roles in activating the host transcriptional response to infection. Like FACT, ETS transcription factors are ancient eukaryotic factors, having originated ~600 million years ago in invertebrate metazoans^29^. Thus, the FEAR pathway appears to both predate and be distinct from, the IFN response. However, only recently was ETS-1 implicated in the host response to microbial infection in bacterial studies where its expression was shown to induced by infection to activate proinflammatory gene expression^30^. Our data indicate that ETS-1 also restricts virus replication and is induced by virus infection in an hSpt16^SUMO^/FACT-dependent manner. Thus, ETS-1 appears to play a broad antimicrobial role in eukaryotes. Further studies are needed to identify genes regulated by ETS-1 during viral infection and their roles in virus restriction. Understanding how FACT is both activated, and potentially countered, by RNA viruses will also be important to determine.

Notably, ETS-1 is a proto-oncoprotein and its upregulation is associated with cancer cell invasiveness and poor survival of cancer patients^36, 37^. Thus, our finding that ETS-1 induction requires hSpt16^SUMO^ function may yield insights into how FACT overexpression promotes oncogenic gene expression and poor outcomes in patients afflicted with malignancies^12, 13^. Thus, our study highlights how characterizing novel viral immune evasion proteins can not only uncover new host antiviral pathways and mechanisms of their regulation, but may also provide tools for probing the function of these pathways in other human etiologies.

## Methods

Specific details regarding the source of all key experimental reagents (primers, plasmids, Abs, cell lines, etc.) can be found in **Supplementary Table 2**.

### Cell lines and primary cultures

Mammalian cell lines were maintained at 37°C in 5% CO_2_ atmosphere. A549, U2OS, HEK293T, HeLa, L929, SIRC, R06E, and BHK-21 cells were cultured in DMEM supplemented with 10% FB Essence (FBE) (Avantor Seradigm). BSC-40 cells were passaged in MEM containing 5% FBE. All media additionally contained 1% non-essential amino acids, 1% L-glutamine, and 1% antibiotic/antimycotic (Gibco). Primary NHDF were passaged in fibroblast growth medium (ATCC) supplemented by Fibroblast Growth Kit with low serum (ATCC). LD652 and Sf21 cells were cultured as previously described at 27°C under normal atmospheric conditions^8, 40^.

### Viruses

Stock preparation and culture of WT and recombinant VV and VSV-GFP^8^ stocks were performed as described^8^. The rescue of a Flag-tagged *A51R* gene encoding alanine at positions 158-162 (ΔA51R^FA51R158-162Ala^) was constructed as described for the ΔA51R^FA51R^ strain^8^ and sequence confirmed by Sanger sequencing of the *A51R* locus. Stocks of YFV-17D-Venus^38^ were obtained from Dr. John Schoggins, and virus particles were collected from culture supernatants after amplification in BHK-21 cells under low MOI conditions. All virus stocks were titrated by plaque assay (VV and VSV-GFP) or fluorescent foci assay (YFV-17D-Venus) on BSC-40 monolayers, with a 1.5% low-melting point agarose (Invitrogen) overlay used for RNA viruses^8^.

Experimental viral infections were incubated for 1 h in serum free DMEM at 37°C before the inoculum was replaced with complete media for the remainder of the infection. Where indicated, virus particles were heat-inactivated prior to infection as described^8^. Where indicated, replacement media after 1 h of infection contained 200 μg/ml AraC (Sigma) and was kept in the media until protein extraction^8^. Lentivirus production in 293T cells via three plasmid transient transfection followed protocols described previously^41^.

### Virus yield assays

For VV: at indicated times post-infection infected cell cultures (supernatants and cells) were both collected by scraping cells into the media, subjected to three rounds of freeze-thaw to release intracellular virus particles and titers were determined by plaque assay on 6-well dishes containing BSC-40 cell monolayers. For VSV and YFV: at indicated times post-infection infected cell culture supernatants were collected, clarified by centrifugation, and clarified supernatant titers were determined by plaque assay (VSV-GFP) or fluorescent foci assay (YFV-17D-Venus) on 6-well dishes containing BSC-40 cell monolayers containing a 1.5% low-melting point agarose overlay.

### Mouse experiments

Six-week-old male BALB/c mice were obtained from the UFMG central animal facility (Belo Horizonte, Brazil) and were kept in ventilated cages with food and water ad libitum. Animals were anaesthetized by intra-peritoneal injection of ketamine and xylazine (70 mg/kg and 12 mg/kg of body weight in PBS, respectively). The intranasal route was used to inoculate 10 μL of PBS or 10 μL of purified viruses (100 PFU) diluted in PBS. Animals were monitored daily for body weight and survival over 15 days and were euthanized if more than 30% of their initial body weight was lost^42^.

### Ethics approval

All mouse model studies were approved by the Committee of Ethics for Animal Experimentation (CETEA) from UFMG, under permission 9/2019.

### Expression vectors

Flag-GFP, VV Flag-A51R, ECTV Flag-A51R, CPXV Flag-A51R, and YLDV Flag-A51R pcDNA3 vectors have been described^8^. The Mpox Flag-A51R gene with flanking *Sac*II and *Pac*I sites was synthesized by Gene Universal and cloned into pCDNA3 using *Sac*II/*Pac*I. N-terminal GFP-Flag-tagged A51R expression vectors were constructed by cloning a *Sac*II/*No*tI fragment from FA51R pCDNA3 into pGFP-C3. GFP-A51R truncations were created by PCR amplification of fragments from A51R templates using iProof DNA Polymerase (Bio-Rad) and cloned into the pGFP-C3 vector. The mutant FA51R encoding alanine substitutions at residues 158-162 was synthesized (Gene Universal) and cloned into pCDNA3 using *Sac*II/*Pac*I cut sites.

Full length HA-hSpt16 pEZ-M06 vector was from GeneCopoeia. HA-hSpt16 mutant fragments (e.g. Δ758-893, 535-554^Ala^, 549-554^Ala^, all K-to-A fragments, etc.) were synthesized (Gene Universal) and subcloned back into the pEZ-M06 vector using appropriate cut sites. GFP-hSpt16 pReceiver lentivirus constructs were purchased from GeneCopoeia. Mutant hSpt16 constructs were first generated in hSpt16-pEZ-M06 (expressing HA-hSpt16) for characterization, and then cloned into pReceiver using appropriate cut sites for lentivirus production and subsequent transduction. All newly amplified products and cloned genes were sequence verified.

### Yeast two-hybrid screen

The coding sequence for VV A51R was PCR-amplified from VV Flag-A51R pCDNA3 and cloned into pB66 as a C-terminal fusion to the Gal4 DNA-binding domain creating Gal4-A51R pB66. The construct was checked by sequencing the entire insert and used as a bait to screen a random-primed Human Lung Cancer cDNA library constructed into pP6 (Hybrigenics Services, Paris, France). pB66 derives from the original pAS2ΔΔ vector ^43^ and pP6 is based on the pGADGH plasmid^44^. 35 million clones (4-fold the complexity of the library) were screened using mating approach with YHGX13 (Y187 ade2-101::loxP-kanMX-loxP, mata) and CG1945 (mata) yeast strains as previously described^43^. His+ colonies were selected on a medium lacking tryptophan, leucine and histidine. The prey fragments of the positive clones were amplified by PCR and sequenced at their 5’ and 3’ junctions. The resulting sequences were used to identify the corresponding interacting proteins in the GenBank database (NCBI).

### Cell line generation

U2OS cell stably expressing full length Flag-hSpt16, GFP-hSpt16 or derivative mutants (ΔNLS or I554A) were generated by lentiviral transduction, followed by puromycin (1.5 μg/ml) selection. Generation of A549 cells stably expressing either control (pLKO.1) or hSpt16-targeted shRNAs were also produced by lentiviral transduction with puromycin (2 μg/ml) selection. Newly generated cell lines underwent at least three rounds of selection prior to use in experiments.

### RNAi and viral infection in cell culture

All siRNAs were obtained from Sigma’s pre-designed siRNA library (see **Supplementary Table 2** for specific siRNA reagents used). Transient siRNA-mediated knockdown was achieved by reverse transfecting A549 or U2OS cells with Lipofectamine 2000 according to manufacturer’s protocol for 48-72 h prior to viral infection.

### Antibodies

Information regarding antibodies used in this study, including their source, is available in **Supplementary Table 2**.

### Immunofluorescence

For staining U2OS GFP-hSpt16 expression cell lines under mock, ΔA51R^FA51R^, or ΔA51R infection conditions, cells were seeded at a density of 30,000 cells per coverslip, cultured overnight, then infected (MOI=3) for 18 hpi. Cells were fixed with methanol, incubated with blocking buffer (PBS with 1% BSA and 0.1% Triton-X) for 1 h, stained with rabbit anti-Flag Ab (Sigma) for 2 h, then incubated with Alexa Fluor-conjugated secondary Ab for 1h. Coverslips were mounted onto glass slides using ProLong™ Diamond anti-fade with DAPI (Thermo Scientific) and imaged on an Olympus Fv10i confocal laser scanning microscope equipped with Fluoview (v.4.2a) and CellSens (v.1.18) software. Similar methods were used for U2OS cells transiently transfected with HA-hSpt16 expression plasmids, with rabbit anti-HA primary Ab (Sigma) used for staining instead.

### General protein extraction

Cells were washed with PBS prior to scraping and transfer into a 1.5 ml microcentrifuge tube for centrifugation at 800 *x g* at 4°C for 15 minutes. Cell pellets were resuspended in either 1x Reporter Lysis Buffer (Promega) containing cOmplete™ EDTA-free Protease Inhibitor Cocktail (Roche) and 1 mM phenylmethylsulfonyl fluoride (PMSF) and freeze-thawed prior to addition of 5x SDS-PAGE loading buffer (100 mM tris HCl, pH 6.8, 4% SDS, 12% (v/v) glycerol, 4 mM DTT, 0.02% (w/v) Bromophenol Blue) or were resuspended in Pierce RIPA buffer (containing cOmplete™ EDTA-free Protease Inhibitor Cocktail and 1 mM PMSF) prior to addition of 5x SDS-PAGE loading buffer. Where indicated, cells were treated with 10 µM TA for 2-4 h prior to protein harvest.

### Immunoprecipitation

For transfection-based Co-IPs, 1×10^6^ 293T cells were seeded prior to transfecting 5 μg of plasmid DNA using Lipofectamine 2000 (Invitrogen), according to manufacturer protocols, in OptiMEM I (Gibco) overnight. Cells were incubated for an additional 24 h after being replaced with complete media. Infection-based Co-IPs involved 4×10^6^ A549 or U2OS cells being seeded overnight, then infected with ΔA51R^FA51R^ (MOI=10). Cells were harvested 18-24 hpi. Prior to cell lysis, cells were washed with equal volumes of PBS twice. Cells were lysed in IP Lysis Buffer (IPLB) (cOmplete™ EDTA-free Protease Inhibitor Cocktail, 1 mM PMSF, 25 mM Tris-HCl at pH 7.4, 150 mM NaCl, 0.5% NP-40) and subjected to shearing and sonication (two-15 second sonications with a 30 second interval). Samples were benzonase (Sigma) treated (250 units/ml) for 1 h at room temperature. 10% of “Input” was collected, and remaining lysate was end-over-end incubated with 5 µg of primary Ab overnight at 4°C [Rabbit-anti-HA (Sigma), Rabbit-anti-Flag (Sigma), Mouse-anti-Spt16 (Biolegend), Mouse-anti-SSRP1 (Biolegend), Mouse-anti-GFP (Biolegend), Mouse-anti-H2BK120ub (Active Motif). Then, lysates were incubated with PureProteome protein A/G magnetic beads (Sigma) for 1-2 h, extensively washed, and immunoprecipitants eluted in 60 μl 2x SDS-PAGE loading buffer. IP of total SUMOylated protein fractions used either SUMO-1 or SUMO-2/3 Ab included with the Signal Seeker SUMOylation 1 or 2/3 Detection Kit (Cytoskeleton) and IPs were carried out according to the manufacturer instructions. Where indicated, nuclear isolations were processed using the Detergent Free Nuclei Isolation Kit (Invent Biotechnologies) according to manufacturer protocols prior to immunoprecipitation.

### Cell fractionation

Cells were infected (MOI=10) for the indicated times prior to fractionation using the Subcellular Protein Fractionation Kit (Thermo). Where indicated, densitometry-based band quantifications were performed using ImageJ software (NIH, v. 1.51n). It is important to note that because the lysis buffer volume used in the fractionation procedure is dependent upon the initial cell pellet size (which varied between infection conditions), and the fractionation buffers were not compatible with protein quantification assays, we only compared cytosolic versus nuclear distribution of a particular protein within (and not between) each infection condition.

### Immunoblotting

Protein extracts were boiled for 10 min prior to SDS-PAGE electrophoresis at 50 V for approximately 4 h. Separated proteins were transferred in Towbin Buffer (BioRad) onto nitrocellulose membranes at 150 mA at 4°C for 90 min and blocked with Odyssey Blocking Buffer (LI-COR) for 1 h at room temperature. Membranes were blotted with primary Ab overnight at 4°C, with actin serving as a loading control. After 3 x 5 min PBS-T (PBS, 0.1% Tween) washes, membranes were incubated in secondary Ab conjugated to an IRDye (LI-COR) for 1 h, washed 3 x 5 min in PBS-T, then a final 5 min PBS wash. Membranes were then imaged with an Odyssey Fc Imager (LI-COR).

### Protein purification

His-tagged A51R (and mutants thereof) were PCR amplified from FA51R pcDNA3 templates using primers encoding an N-terminal His tag and *Nco*I/*Not*I cut sites used for cloning into the pET22b bacterial expression vector. His-A51R pET22b vectors were transformed into BL21(DE3) *E. coli* cells then grown in Luria broth at 37°C, induced at mid-log phase with IPTG (0.5 mM), then harvested by centrifugation (5 krpm, 20 min, 4°C) after 4 h prior to lysis by sonication in binding buffer (50 mM Tris-HCl pH 7.4, 500 mM NaCl, 10% glycerol, 50 mM imidazole, 4°C). Soluble fractions were obtained by centrifugation, supernatant loaded onto a HisTrap HP 1 ml column (GE Helthcare), washed, and eluted with elution buffer (50 mM Tris-HCl pH 7.4, 500 mM NaCl, 500 mM imidazole, 10% glycerol). Pooled His-A51R factions were run over a Superdex 200 Increase 10/300 size-exclusion chromatography (SEC) column (GE Healthcare), exchanged into Storage Buffer (50 mM Tris-HCl pH 7.4, 500 mM NaCl, 10% glycerol) and concentrated prior to storage. His-tagged mCherry (mCherry-His) was PCR amplified from pmCherry-C1 vector (Takara) templates with primers encoding a His-tag and *Nco*I/*Not*I cut sites (Takara), cloned into pET22b, and expressed and purified using HisTrap HP 1ml columns as described above.

Flag-hSpt16 proteins were purified from HeLa cells transduced with Flag-hSpt16-expressing lentivirus following protocols previously described^45^ with minor modifications. Briefly, cells were grown to confluency and harvested in PBS prior to centrifugation at 1,200 *x g*, 15 min, 4°C. Cells were subsequently lysed in Buffer A (20 mM Tris pH7.5, 0.2 M NaCl, 5% glycerol, 0.01 mM Octyl-beta-D-thioglucopyranoside, 2 mM DTT, 250 units/ml benzonase) via sonication. Lysates were subjected to centrifugation at 18,000 rpm for 15 min, 0.22 µm filtered, then poured onto loaded onto a pre-equilibrated HiTrap DEAE FF 5 ml column (GE Healthcare) into Buffer B (20 mM Tris pH 7.5, 1.0 M NaCl, 5% glycerol, 0.01 mM Octyl-beta-D-thioglucopyranoside, 2 mM DTT). Flag-hSpt16 containing fractions were combined and incubated with M2 resin (Sigma) overnight. M2 resin was washed using a gravity flow column (Bio-Rad) then competed off with Flag peptides (0.5 mg/ml) for 30 min. Samples were eluted in M2 buffer and Flag-Spt16 containing fractions were pooled, concentrated using Amicon centrifugal filter unit (50 kDa cut-off, Millipore), aliquoted, snap frozen in liquid nitrogen and stored at 80°C. Where indicated, Flag-hSpt16 proteins were incubated with 12.5 U of recombinant ULP-1 SUMO Protease (Sigma) for 16 h at 4°C to remove SUMO moieties.

His-tagged SSRP1 was PCR amplified from SSRP1 pReceiver (GeneCopoeia) using primers encoding an N-terminal His tag and *Nde*I/*Not*I cut sites for cloning into pET22b. His-SSRP1 pET22b was used to transform BL21(DE3) cells that were grown at 37°C in standard Luria-Bertani medium plus appropriate antibiotics. Expression was induced by the addition of IPTG to a final concentration of 0.5 mM at OD 600 nm = 0.5 AU. Cultures were incubated for 4 h after induction and the cells were harvested by centrifugation (5,000 rpm, 20 min, 4°C). The bacterial pellet was resuspended in binding buffer (50 mM Tris-HCl pH 7.4, 500 mM NaCl, 10% glycerol, 50 mM imidazole) at 4°C. Cell lysis was carried out by sonication and the soluble fraction was obtained by centrifugation (10,000xg, 30 min, 4°C).

The supernatant was filtered with a 0.22 μm filter and was loaded onto a pre-equilibrated HisTrap HP 1 ml column (GE Healthcare) at 0.5 ml/min with a peristaltic pump at 4°C. The column was washed with 10 column volumes of binding buffer and the column was eluted by 5 column volumes of elution buffer (50 mM Tris-HCl pH 7.4, 500 mM NaCl, 500 mM imidazole, 10% glycerol). The fractions containing His-SSRP1 were concentrated with an Amicon centrifugal filter unit (50 kDa cut-off, Millipore) and buffer exchanged into storage buffer (50 mM Tris-HCl pH 7.4, 500 mM NaCl, 10% glycerol). The protein was aliquoted, snap frozen in liquid nitrogen and stored at −80°C. The co-expressed/purified hSpt16 and SSRP1 proteins for use in FACT-nucleosome binding assays were expressed using a baculovirus expression system in Sf21 cells and purified as described^45^.

### *in vitro* Co-IPs and pulldowns

For histone pulldown assays, 5 μg of His-tagged H2A:H2B or His-H3:Flag-H4 (Diagenode) and 2 μg of Flag-hSpt16 were added to 250 μl of binding buffer (50 mM Tris-HCl pH 7.4, 150 mM NaCl, 0.1% NP-40, 50 mM imidazole) and the mixture was end-over-end mixed at 4°C overnight. The next day, 10% input aliquots were taken and the rest of the sample was incubated with HIS-Select® Nickel Magnetic Agarose Beads (Sigma) for 1 h, extensively washed with binding buffer, and eluted in 60 μl 2x SDS-PAGE loading buffer. For His-A51R or His-SSRP1 nickel bead pulldowns, 1.6 µg His-A51R or 2.3 µg His-SSRP1 were added to 500 µl binding buffer containing 1.5 µg Flag-hSpt16 and the pulldown was performed as described above. For Flag-hSpt16 Co-IPs, 1.6 µg His-A51R or 2.3 µg His-SSRP1 were added to 500 µl of IPLB containing 1.5 µg Flag-hSpt16, and the mixture was end-over-end mixed overnight at 4°C. The next day, 10% input aliquots were taken and the rest of the sample was incubated with 5 µg of Flag Ab for 2 h, and then incubated with PureProteome protein A/G magnetic beads (Sigma) for 1 h, extensively washed, and immunoprecipitants were eluted 60 μl 2x SDS-PAGE loading buffer. For in vitro competition assays, His-A51R and Flag-hSpt16 or His-SSRP1 and Flag-hSpt16 proteins were incubated in binding buffer overnight to form complexes prior to addition of indicated amounts of either His-SSRP1 or His-A51R for 3 h. After this incubation period, 5 µg of SSRP1 or Flag Ab was added for 2 h, PureProteome protein A/G magnetic beads (Sigma) then added for 1 h, extensively washed, and immunoprecipitants were eluted in 60 μl 2x SDS-PAGE loading buffer. A similar procedure was used for Flag-hSpt16-Tubulin Co-IP assays performed in the absence/presence of His-A51R but, where indicated, 5 µg of purified porcine tubulin (Cytoskeleton) was present in protein mixtures prior to Flag Ab immunoprecipitation.

### Nucleosome complex reconstitution & binding assays

Human recombinant histones H2A, H2B, H3 and H4 were expressed, refolded and purified as described^46^. To visualize the histone components, Atto 647N–labeled H2A–H2B and Alexa 488–labeled H3–H4 were utilized in the FACT binding assay as described ^45, 47^. Briefly, human FACT was pre-mixed with refolded H2A-H2B at equimolar for 10 min at room temperature, followed by adding an equimolar of either (H3-H4)_2_ tetrasome reconstituted onto 79 bp Widom 601 DNA or (H2A-H2B)-(H3-H4)_2_ hexasome reconstituted onto 128 bp Widom 601 DNA. The binding assay was performed in 20 mM Tris-Cl, pH 7.5, 150 mM NaCl, 1 mM EDTA and 1 mM TCEP. After a 30 min incubation, the complex was visualized in 5% native PAGE by Typhoon scan. Where indicated, FACT complexes were de-SUMOylated with ULP-1 SUMO Protease (Sigma) prior to mixing with tetrasomes or hexasomes.

### Microtubule co-sedimentation assays

The Microtubule co-sedimentation assay was performed as described^48^ with minor modifications. Briefly, 40 μM of porcine brain tubulin (Cytoskeleton Inc.) in 80 mM PIPES pH 6.9, 2 mM MgCl_2_, 0.5 mM EGTA, 5% glycerol, 1 mM GTP was polymerized into MTs by incubation at 37°C for 20 min. MTs were then stabilized by resuspending in 50 mM Tris-HCl, pH 6.8, 10% glycerol buffer supplemented with 100 µM paclitaxel (Sigma). A constant amount of MT was incubated alone or with either equal concentrations of His-A51R, its mutants or mCherry-His protein as a negative control. The samples were incubated for 30 min at 37°C, then were layered over 30 μl of a 45% glycerol cushion (50 mM Tris-HCl pH 7.4, 100 mM NaCl, 100 μM paclitaxel, 45% glycerol) and were subsequently spun at 16,000 x g for 30 min at 37°C to pellet MTs and bound proteins. Supernatant and pellet fractions were separated and resuspended in 2x SDS-PAGE loading buffer, and equal amounts of supernatant and pellet were run on 10% Tris-HCl stain-free gels (Bio-Rad). Gels were imaged using Bio-Rad ChemiDoc^TM^ Touch Imaging System equipped with Image Lab Software (v. 6.1).

## RNA-Sequencing

### RNA extraction

A549 cells expressing control or hSpt16 shRNA were mock-infected or infected with ΔA51R^FA51R^ or ΔA51R strains for 1 h (MOI=50). Following infection, complete media was added and incubated until 4 hpi when cells were lysed, and total RNA was extracted using the RNeasy Mini kit (Qiagen) according to manufacturer’s protocol.

### Library Preparation and Sequencing

RNA samples were quantified using Qubit 2.0 Fluorometer (Life Technologies, Carlsbad, CA, USA) and RNA integrity was checked using Agilent TapeStation 4200 (Agilent Technologies, Palo Alto, CA, USA). rRNA depletion was performed using Ribozero rRNA Removal Kit (Illumina, San Diego, CA, USA). RNA sequencing library preparation used NEBNext Ultra RNA Library Prep Kit for Illumina by following the manufacturer’s recommendations (NEB, Ipswich, MA, USA). Sequencing libraries were validated using the Agilent Tapestation 4200 (Agilent Technologies, Palo Alto, CA, USA), and quantified by using Qubit 2.0 Fluorometer (Invitrogen, Carlsbad, CA) as well as by quantitative PCR (Applied Biosystems, Carlsbad, CA, USA). The sequencing libraries were clustered on two lanes of a flowcell. After clustering, the flowcell was loaded on the Illumina HiSeq instrument (4000 or equivalent) according to manufacturer’s instructions. The samples were sequenced using a 2×150bp Paired End (PE) configuration. Image analysis and base calling were conducted by the HiSeq Control Software (HCS). Raw sequence data (.bcl files) generated from Illumina HiSeq was converted into fastq files and de-multiplexed using Illumina’s bcl2fastq 2.17 software. One mismatch was allowed for index sequence identification.

### Data Analysis

Quality of the raw FASTQ files were checked using FASTQC (https://www.bioinformatics.babraham.ac.uk/projects/fastqc/). Trimmomatic (v.0.36) removed possible adapter sequences and nucleotides with poor quality^49^. Trimmed reads were mapped to the Human (hg38/GRCh38) using the STAR aligner (v.2.5.2b)^50^. Using the aligned BAM files, unique gene hit counts were calculated by using featureCounts^51^. Following extraction of gene hit counts, DESeq2 (v.1.20) was used for differential expression analysis by comparing gene expression between the sample groups^52^. The Wald test was used to generate p-values and log2 fold changes. Genes with adjusted p-values <0.05 and absolute log2 fold changes >1 were termed differentially expressed genes for each comparison then further curated using an adjusted p-value <0.01 and absolute log2 fold change >0.58 to generate a DEG list. We then identified overlapping genes across multiple conditions at 4 hpi and generated a master list of hSpt16-regulated genes by combining comparisons where differential gene expression was dependent only on hSpt16 expression regardless of the presence of infection. This list was overlapped with conditions where differential expression was dependent on A51R. The resulting gene list indicates DEGs that were dependent on hSpt16 and A51R (**Fig. 5e, Supplementary Table 1g**). The overlapping DEGs in **Fig. 5e** that were down-regulated after hSpt16 RNAi and down-regulated in the presence of A51R were further analyzed for immune system related function. Genes were curated as “immunity genes” if they were found to belong to the “immune system process (GO: 0002376)” or “response to stimulus (GO: 0050896) gene ontology groups using PANTHER^53^ (v.17.0) or QuickGO^54^ or if evidence was found for immune system involvement through manual literature research (**Supplementary Table 1f**). This list was then interrogated for genes encoding known TFs. The relative expression of 8 immunity-related TF genes identified among this gene list were then displayed in heat map-based presentations of these data (**Fig. 5f**) using Graphpad Prism (v.8.0) software.

### ChIP-PCR

For each treatment, 15 x 10^6^ A549 cells were subjected to ChIP using Go-ChIP-Grade anti-Spt16 mouse Ab or isotype control Ab (BioLegend) and ChIP-IT® Express Chromatin Immunoprecipitation Kits (Active Motif) protocols according to manufacturer protocols. Input and immunoprecipitated DNA was then subjected to PCR reactions containing iProof DNA Polymerase (Bio-Rad) and primers targeting a promoter-proximal region of the *ets-1* gene (chr11: nts 128,391,556-128,392,408) previously identified in hSpt16 ChIP-seq studies^31^.

### Statistical analyses

All statistical analyses were conducted using GraphPad Prism (v.8.0) software and *P* values <0.05 were considered statistically significant. Sample sizes, statistical tests used, and *P* values (only those <0.05, indicated with horizontal brackets of treatments being compared) are indicated in the respective figure legend or figure for each quantitative experiment. *P* values <0.0001 are indicated as “*P*<0.0001” rather than with exact *P* values.

## Biological materials

Plasmids, primers, strains, and any other research reagents generated by the authors will be distributed upon request to other research investigators under a Material Transfer Agreement.

## Data availability

The authors declare that the main data supporting the findings of this study are available within the article and its Supplementary Information. RNA-seq data sets are available under Geo accession: GSE185829. Any additional information required to reanalyze the data reported in this paper is available from the corresponding author upon request.

## Supplementary Information

**Supplementary Table 1**| RNA-seq analyses 4 hpi in mock-, ΔA51R^FA51R^, and ΔA51R-infected control shRNA- or hSpt16 shRNA-expressing A549 cells.

1a-Guide to Supplementary Table 1.

1b-DEGs in mock-infected cells after hSpt16 RNAi.

1c-DEGs in ΔA51R^FA51R^-infected cells after hSpt16 RNAi. 1d-DEGs in ΔA51R-infected cells after hSpt16 RNAi.

1e-DEGs after hSpt16 RNAi in either mock or VV-infected conditions.

1f-DEGs between ΔA51R^FA51R^ and ΔA51R under control RNAi conditions.

1g-DEGs from hSpt16 RNAi experiments that are also DEGs between ΔA51R^FA51R^ and ΔA51R infections.

1h-Immunity-related DEGS from Table S1G that are down-regulated in one or more hSpt16 RNAi treatments and that are also down-regulated between ΔA51R^FA51R^ and ΔA51R infections.

**Supplementary Table 2**| Key Experimental Reagents.

## Acknowledgements

We thank colleagues Drs. Ivan D’Orso, Julie Pfeiffer and Nick Conrad (UTSW) and Dr. Duane Winkler (UT Dallas) for helpful discussions. This work was supported by grants to DBG from the NIH (1R21AI144203-01 and 1R35GM137978-01) and Welch Foundation (I-2062-20210327) and by funding to DBG from the UTSW Endowed Scholars Program. CP was supported by NIH Training Grant T32 AI007520. Animal experiments were supported by funding to FG by the INCTV Initiative.

## Author contributions

Conceptualization, D.B.G. and E.A.R.; Methodology, D.B.G., E.A.R.; Investigation, D.B.G., E.A.R, D.S., S.C., C.P., S.B.O., A.E., D.H., Y.L., R.O.; K.L., O.V.H., N.K.C.; Data Curation, D.B.G. and SC.; Writing—original draft, D.B.G. Writing—Review & Editing, K.L., N.M.A.,, R.O., and F.G.; Visualization, D.B.G. and E.A.R.; Supervision, D.B.G., K.L., N.M.A.,, R.O., and F.G.; Funding Acquisition, D.B.G., C.P., and F.G.

## Additional information

**Correspondence and request for materials** should be addressed to Don Gammon.

## Extended Data

**Extended Data Fig. 1.**
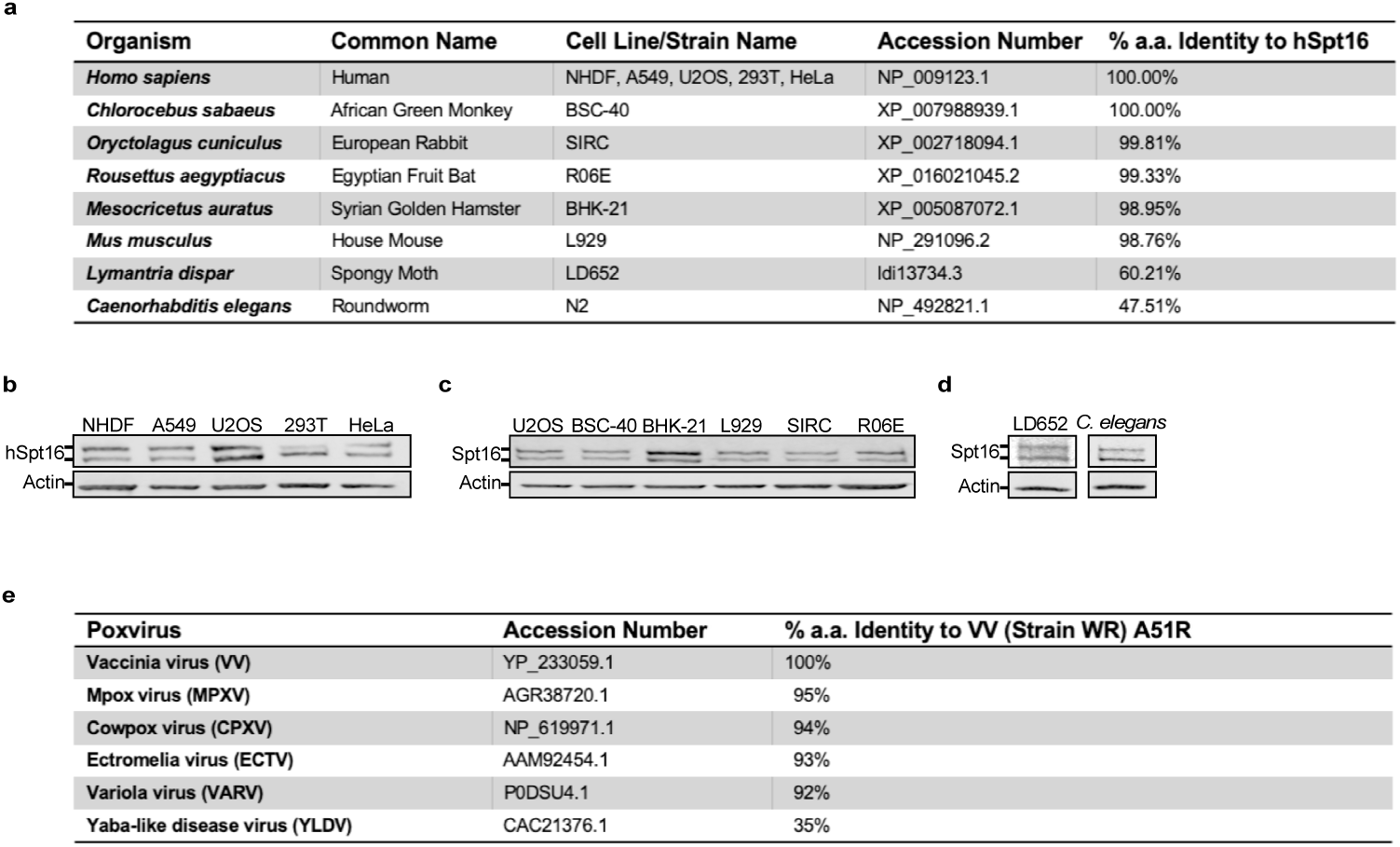
Conservation of Spt16, Spt16 SUMOylation, and poxvirus A51R proteins. **a**, Amino acid (a.a.) similarity of eukaryotic Spt16 proteins. **b**-**d**, IB of endogenous Spt16 proteins from indicated human (**b**) and mammalian animal cell lines (**c**) and from invertebrate cell lines and animal tissues (*C. elegans*) (**d**) showing SUMOylated and non-SUMOylated Spt16 forms. Non-human cell lines derive from the following: BSC-40 (*Chlorocebus sabaeus*); BHK-21 (*Mesocricetus auratus*); L929 (*Mus musculus*); SIRC (*Oryctolagus cuniculus*); R06E (*Rousettus aegyptiacus*); and LD652 (*Lymantria dispar*). **e**, Overall amino acid similarity of indicated poxvirus A51R proteins. Images in **b**-**d**, are representative of at least two independent experiments.

**Extended Data Fig. 2.**
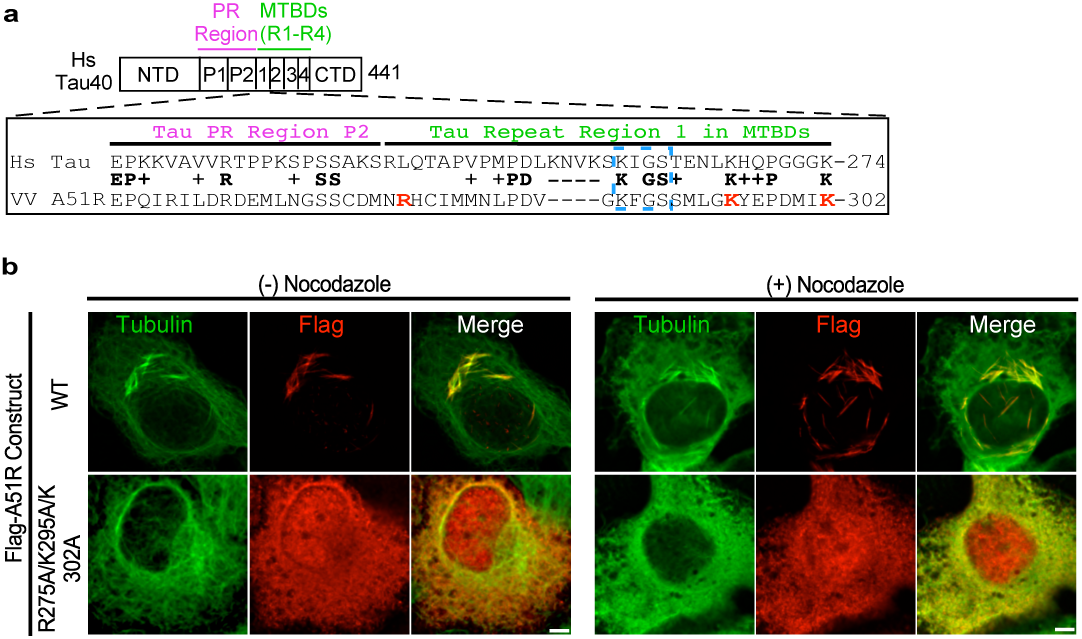
A Tau-like motif in VV A51R mediates MT interaction. **a**, Alignment of VV A51R with Proline Rich (PR) Region and Region 1 of human Tau40 MT-binding domains (MTBD)^55^. “KXGS” motif conserved in all four MTBDs in Tau is boxed. Residues in red indicate those converted to alanine in the triple A51R mutant. **b**, IF staining of U2OS cells transfected with WT or triple mutant FA51R expression constructs after 24 h. Where indicated, 40 μM nocodazole was added to depolymerize MTs 6 h post-transfection MTs. Scale bar, 5 μm. Images in **a**,**b** are representative of at least two independent experiments.

**Extended Data Fig. 3.**
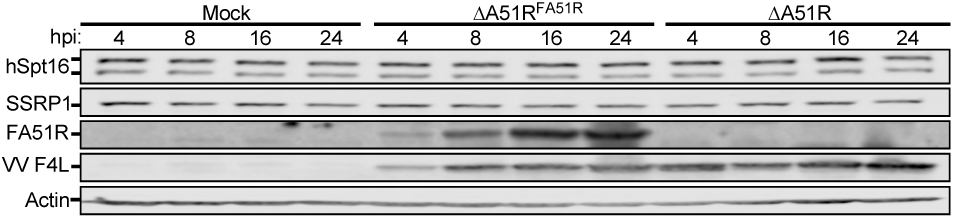
VV A51R does not affect total hSpt16 protein levels. IB of hSpt16 in U2OS WCE at indicated times post-infection in U2OS WCE (MOI=3). Image is representative of two independent experiments.

**Extended Data Fig. 4.**
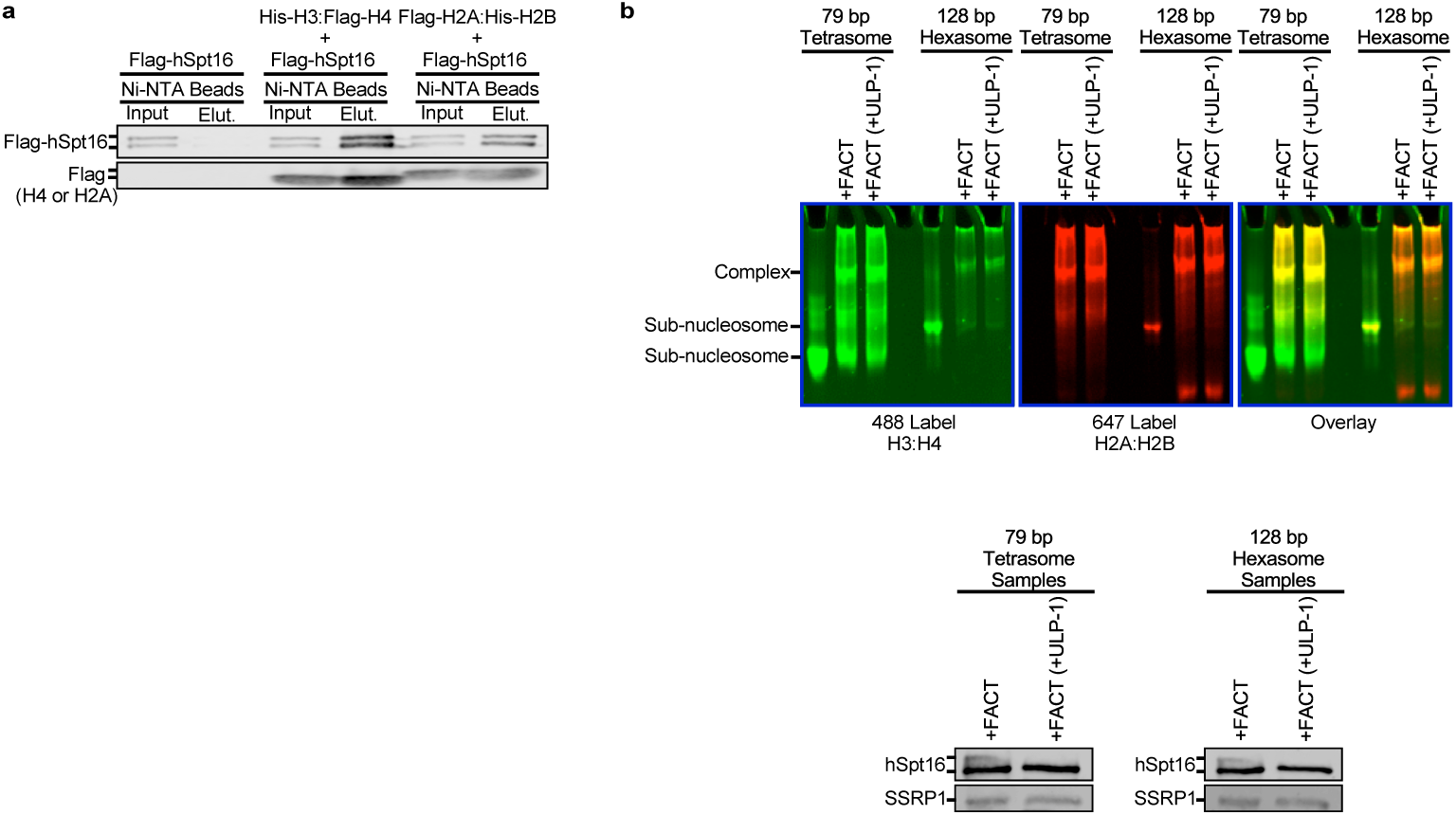
hSpt16 SUMOylation does not affect histone or nucleosome interactions *in vitro*. **a**, *in vitro* nickel bead pulldown of His-tagged H3/H4 and H2A/H2B complexes with purified Flag-hSpt16. **b**, *in vitro* FACT binding assay with reconstituted (H3-H4)_2_ tetrasomes or (H2A-H2B)-(H3-H4)_2_ hexasomes^45^. Where indicated, purified FACT complexes were treated with ULP-1 to remove hSpt16 SUMOylation prior to being added to binding assays. IBs of FACT treatments are shown below indicating hSpt16 and SSRP1 levels. Images in **a**,**b** representative of two independent experiments.

**Extended Data Fig. 5.**
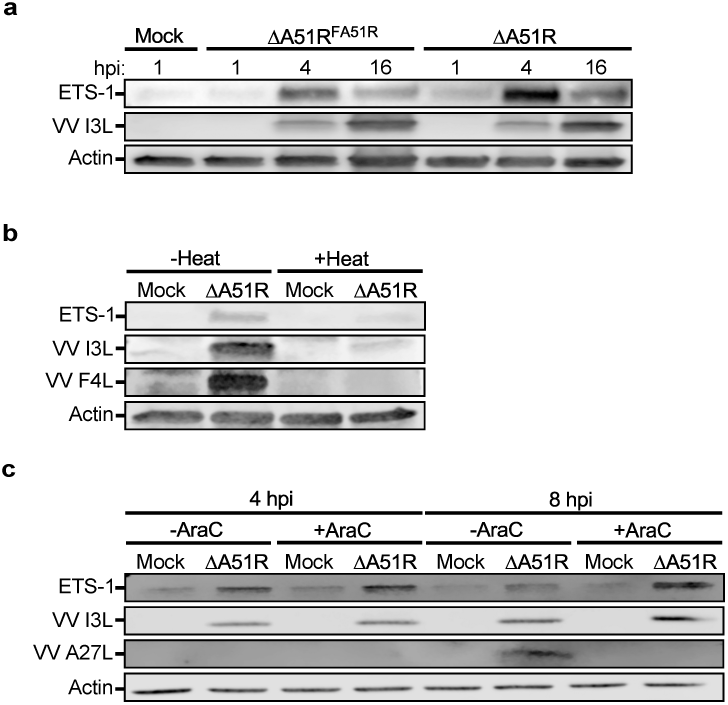
ETS-1 induction by VV infection occurs by 4 hpi and requires early VV gene expression but not viral DNA replication. **a**, IB of ETS-1 in WCE from A549 cells infected with indicated strains for indicated times. **b**, IB of ETS-1 in WCE from A549 cells under indicated infection conditions 4 hpi. Where indicated, inoculum was heat-inactivated prior to infection. **c**, IB of ETS-1 in WCE from A549 cells under indicated infection conditions. Where indicated, AraC was added to media 1 hpi. VV A27L is a late VV protein and its expression requires VV DNA replication. Images in **a-c** are representative of two independent experiments.

**Extended Data Fig. 6.**
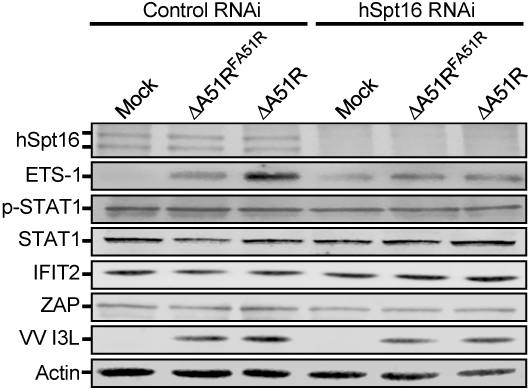
IFN pathway activation in A549 cells is unchanged between ΔA51R^FA51R^ and ΔA51R infections or in the presence of hSpt16 depletion. **a**, IB of activated IFN pathway signaling components (phospho-STAT1) and IFN-stimulated gene (ZAP, IFIT2) expression in WCE of control or hSpt16-shRNA-expressing A549 cell lines 4 hpi with indicated VV strains (MOI=10). Image is representative of two independent experiments.

**Extended Data Fig. 7.**
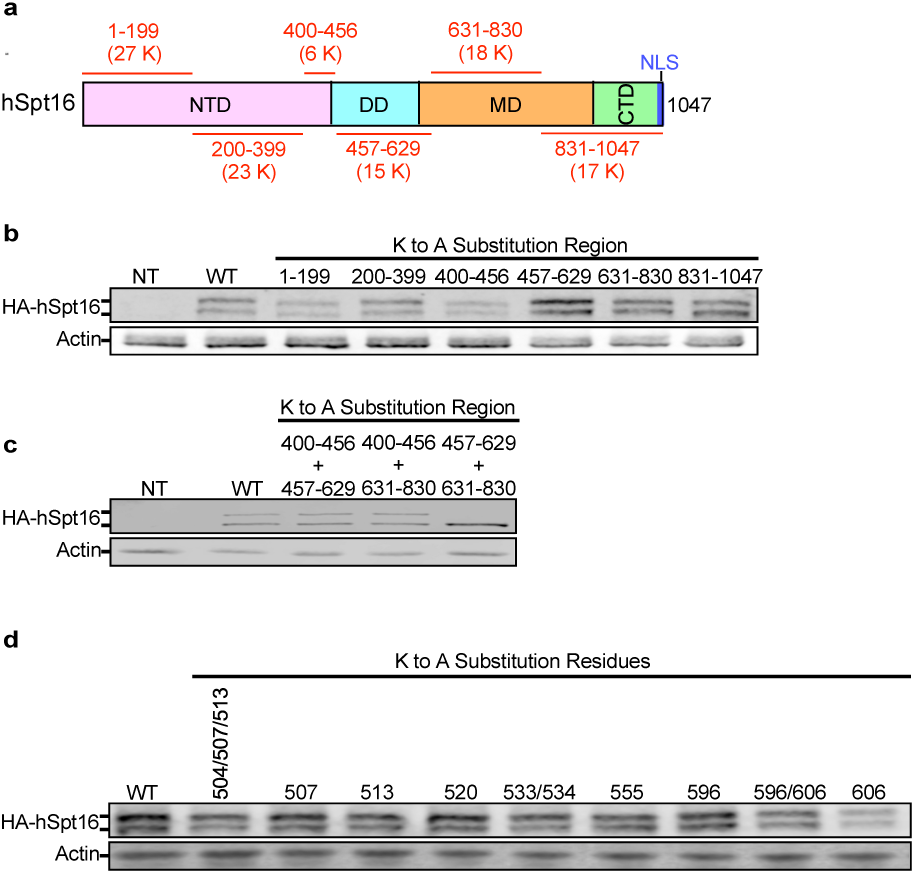
Characterization of hSpt16 SUMOylation site(s). **a**, hSpt16 protein map indicating regions containing K to A substitutions in HA-hSpt16 mutant constructs and the number of K residues in each region. **b**,**c**, IB of WT and mutant HA-hSpt16 transfected constructs in 293T WCE containing K to A substitutions at all K residues within indicated a.a. regions of hSpt16. **d**, IB of WT and mutant HA-hSpt16 transfected constructs in 293T WCE containing K to A substitutions at indicated sites in hSpt16. Images in **b-d** are representative of at least two independent experiments.

## References

1. Nan, Y., Nan, G. & Zhang, Y.J. Interferon induction by RNA viruses and antagonism by viral pathogens. Viruses 6, 4999–5027 (2014).

2. Beachboard, D.C. & Horner, S.M. Innate immune evasion strategies of DNA and RNA viruses. Curr. Opin. Microbiol. 32, 113–119 (2016).

3. McFadden, G. Poxvirus tropism. Nat. Rev. Microbiol. 3, 201–213 (2005).

4. Yang, Z. Monkeypox: A potential global threat? J. Med. Virol. 94, 4034–4036 (2022).

5. Mohapatra, R.K. et al. Unexpected sudden rise of human monkeypox cases in multiple non-endemic countries amid COVID-19 pandemic and salient counteracting strategies: Another potential global threat? Int. J. Surg. 103, 106705 (2022).

6. Yu, H., Bruneau, R.C., Brennan, G. & Rothenburg, S. Battle Royale: Innate Recognition of Poxviruses and Viral Immune Evasion. Biomedicines 9 (2021).

7. Reus, J.B., Rex, E.A. & Gammon, D.B. How to Inhibit Nuclear Factor-Kappa B Signaling: Lessons from Poxviruses. Pathogens 11 (2022).

8. Gammon, D.B. et al. A single vertebrate DNA virus protein disarms invertebrate immunity to RNA virus infection. Elife 3 (2014).

9. Orphanides, G., Wu, W.H., Lane, W.S., Hampsey, M. & Reinberg, D. The chromatin-specific transcription elongation factor FACT comprises human SPT16 and SSRP1 proteins. Nature 400, 284–288 (1999).

10. Orphanides, G., LeRoy, G., Chang, C.H., Luse, D.S. & Reinberg, D. FACT, a factor that facilitates transcript elongation through nucleosomes. Cell 92, 105–116 (1998).

11. LeRoy, G., Orphanides, G., Lane, W.S. & Reinberg, D. Requirement of RSF and FACT for transcription of chromatin templates in vitro. Science 282, 1900–1904 (1998).

12. Garcia, H. et al. Facilitates chromatin transcription complex is an “accelerator” of tumor transformation and potential marker and target of aggressive cancers. Cell Rep 4, 159–173 (2013).

13. Fleyshman, D. et al. Level of FACT defines the transcriptional landscape and aggressive phenotype of breast cancer cells. Oncotarget 8, 20525–20542 (2017).

14. Lolis, A.A. et al. Myogenin recruits the histone chaperone facilitates chromatin transcription (FACT) to promote nucleosome disassembly at muscle-specific genes. J. Biol. Chem. 288, 7676–7687 (2013).

15. Minsky, N. et al. Monoubiquitinated H2B is associated with the transcribed region of highly expressed genes in human cells. Nat. Cell Biol. 10, 483–488 (2008).

16. Safina, A. et al. Complex mutual regulation of facilitates chromatin transcription (FACT) subunits on both mRNA and protein levels in human cells. Cell Cycle 12, 2423–2434 (2013).

17. Schoggins, J.W. et al. A diverse range of gene products are effectors of the type I interferon antiviral response. Nature 472, 481–485 (2011).

18. Gasparian, A.V. et al. Curaxins: anticancer compounds that simultaneously suppress NF-kappaB and activate p53 by targeting FACT. Sci. Transl. Med. 3, 95ra74 (2011).

19. Suzawa, M. et al. A gene-expression screen identifies a non-toxic sumoylation inhibitor that mimics SUMO-less human LRH-1 in liver. Elife 4 (2015).

20. Li, S.J. & Hochstrasser, M. A new protease required for cell-cycle progression in yeast. Nature 398, 246–251 (1999).

21. Winkler, D.D. & Luger, K. The histone chaperone FACT: structural insights and mechanisms for nucleosome reorganization. J. Biol. Chem. 286, 18369–18374 (2011).

22. Dehmelt, L. & Halpain, S. The MAP2/Tau family of microtubule-associated proteins. Genome Biol. 6, 204 (2005).

23. Brameier, M., Krings, A. & MacCallum, R.M. NucPred--predicting nuclear localization of proteins. Bioinformatics 23, 1159–1160 (2007).

24. Kosugi, S., Hasebe, M., Tomita, M. & Yanagawa, H. Systematic identification of cell cycle-dependent yeast nucleocytoplasmic shuttling proteins by prediction of composite motifs. Proc. Natl. Acad. Sci. U. S. A. 106, 10171–10176 (2009).

25. Pavri, R. et al. Histone H2B monoubiquitination functions cooperatively with FACT to regulate elongation by RNA polymerase II. Cell 125, 703–717 (2006).

26. Fleming, A.B., Kao, C.F., Hillyer, C., Pikaart, M. & Osley, M.A. H2B ubiquitylation plays a role in nucleosome dynamics during transcription elongation. Mol. Cell 31, 57–66 (2008).

27. Sanda, C. et al. Differential gene induction by type I and type II interferons and their combination. J Interferon Cytokine Res 26, 462–472 (2006).

28. Goulet, M.L. et al. Systems analysis of a RIG-I agonist inducing broad spectrum inhibition of virus infectivity. PLoS Pathog. 9, e1003298 (2013).

29. Garrett-Sinha, L.A. Review of Ets1 structure, function, and roles in immunity. Cell Mol Life Sci 70, 3375–3390 (2013).

30. Teng, Y. et al. Expression of ETS1 in gastric epithelial cells positively regulate inflammatory response in Helicobacter pylori-associated gastritis. Cell Death Dis. 11, 498 (2020).

31. Kolundzic, E. et al. FACT Sets a Barrier for Cell Fate Reprogramming in Caenorhabditis elegans and Human Cells. Dev. Cell 46, 611–626 e612 (2018).

32. Deng, L. et al. Suppression of NF-kappaB Activity: A Viral Immune Evasion Mechanism. Viruses 10 (2018).

33. Shaw, A.E. et al. Fundamental properties of the mammalian innate immune system revealed by multispecies comparison of type I interferon responses. PLoS Biol. 15, e2004086 (2017).

34. Rahman, M.M. & McFadden, G. Modulation of NF-kappaB signalling by microbial pathogens. Nat. Rev. Microbiol. 9, 291–306 (2011).

35. Smale, S.T. Transcriptional regulation in the innate immune system. Curr. Opin. Immunol. 24, 51–57 (2012).

36. Fry, E.A. & Inoue, K. Aberrant expression of ETS1 and ETS2 proteins in cancer. Cancer Rep Rev 2 (2018).

37. Dittmer, J. The role of the transcription factor Ets1 in carcinoma. Semin. Cancer Biol. 35, 20–38 (2015).

38. Hanners, N.W. et al. Western Zika Virus in Human Fetal Neural Progenitors Persists Long Term with Partial Cytopathic and Limited Immunogenic Effects. Cell Rep 15, 2315–2322 (2016).

39. Mahajan, R., Delphin, C., Guan, T., Gerace, L. & Melchior, F. A small ubiquitin-related polypeptide involved in targeting RanGAP1 to nuclear pore complex protein RanBP2. Cell 88, 97–107 (1997).

40. Winkler, D.D., Muthurajan, U.M., Hieb, A.R. & Luger, K. Histone chaperone FACT coordinates nucleosome interaction through multiple synergistic binding events. J. Biol. Chem. 286, 41883–41892 (2011).

41. Aboulaich, N. Lentivirus Production. Bio Protoc A Bio-101: e39. (2011).

42. Lourenco, K.L., Chinalia, L.A., Henriques, L.R., Rodrigues, R.A.L. & da Fonseca, F.G. Zoonotic vaccinia virus strains belonging to different genetic clades exhibit immunomodulation abilities that are proportional to their virulence. Virol J. 18, 124 (2021).

43. Fromont-Racine, M., Rain, J.C. & Legrain, P. Toward a functional analysis of the yeast genome through exhaustive two-hybrid screens. Nat. Genet. 16, 277–282 (1997).

44. Bartel, P.L. & Fields, S. Analyzing protein-protein interactions using two-hybrid system. Methods Enzymol. 254, 241–263 (1995).

45. Wang, T. et al. The histone chaperone FACT modulates nucleosome structure by tethering its components. Life Sci Alliance 1, e201800107 (2018).

46. Dyer, P.N. et al. Reconstitution of nucleosome core particles from recombinant histones and DNA. Methods Enzymol. 375, 23–44 (2004).

47. Liu, Y. et al. FACT caught in the act of manipulating the nucleosome. Nature 577, 426–431 (2020).

48. Kesten, C., Schneider, R. and Persson, S. . n vitro Microtubule Binding Assay and Dissociation Constant Estimation. Bio Protoc 6(6): e1759 (2016).

49. Bolger, A.M., Lohse, M. & Usadel, B. Trimmomatic: a flexible trimmer for Illumina sequence data. Bioinformatics 30, 2114–2120 (2014).

50. Dobin, A. et al. STAR: ultrafast universal RNA-seq aligner. *Bioinformatics (Oxford*, England) 29, 15–21 (2013).

51. Liao, Y., Smyth, G.K. & Shi, W. featureCounts: an efficient general purpose program for assigning sequence reads to genomic features. Bioinformatics 30, 923–930 (2013).

52. Love, M.I., Huber, W. & Anders, S. Moderated estimation of fold change and dispersion for RNA-seq data with DESeq2. Genome Biology 15, 550 (2014).

53. Mi, H., Muruganujan, A. & Thomas, P.D. PANTHER in 2013: modeling the evolution of gene function, and other gene attributes, in the context of phylogenetic trees. Nucleic Acids Res. 41, D377–386 (2013).

54. Binns, D. et al. QuickGO: a web-based tool for Gene Ontology searching. Bioinformatics 25, 3045–3046 (2009).

55. Barbier, P. et al. Role of Tau as a Microtubule-Associated Protein: Structural and Functional Aspects. Front. Aging Neurosci. 11, 204 (2019).

