## Supplementary Table 2 for "A FACT-ETS-1 Antiviral Response Pathway Restricts Viral Replication and is Countered by Poxvirus A51R Proteins"

**Supplementary Table 2- Key Experimental Reagents**

| REAGENT or RESOURCE | SOURCE | IDENTIFIER |
|---|---|---|
| Antibodies | | |
| Rabbit polyclonal anti-Flag | Sigma-Aldrich | Cat# F7425, RRID:AB_439687 |
| Mouse monoclonal anti-Flag | Wako | Cat# 014-22383, RRID:AB_10660291 |
| Rabbit polyclonal anti-HA | Sigma-Aldrich | Cat# H6908, RRID:AB_260070 |
| Mouse monoclonal anti-HA.11 | BioLegend | Cat# 901515, RRID:AB_2565334 |
| Mouse monoclonal anti-GFP | BioLegend | Cat# 902601, RRID:AB_2565021 |
| Mouse monoclonal anti-His | BioLegend | Cat# 906102, RRID:AB_2565062 |
| Rabbit monoclonal anti-Spt16 | Cell Signaling Technology | Cat# 12191, RRID:AB_2732025 |
| Mouse monoclonal anti-Spt16 | BioLegend | Cat# 607002, RRID:AB_315689 |
| Mouse monoclonal anti-Spt16 Go-ChIP Grade | BioLegend | Cat# 607008, RRID:AB_2721598 |
| Mouse monoclonal anti-Spt16 | Santa Cruz Biotechnology | Cat# sc-165987, RRID:AB_2286916 |
| Rabbit monoclonal anti-SSRP1 | Cell Signaling Technology | Cat# 13421, RRID:AB_2714160 |
| Mouse monoclonal anti-SSRP1 | BioLegend | Cat# 609702, RRID:AB_315731 |
| Mouse monoclonal anti-SSRP1 | Santa Cruz Biotechnology | Cat# sc-74536, RRID:AB_1129655 |
| Rabbit monoclonal anti-SUMO1 | Abcam | Cat# ab32058, RRID:AB_778173 |
| Mouse monoclonal anti SUMO2+3 | Abcam | Cat# ab81371, RRID:AB_1658424 |
| Mouse monoclonal anti-Histone H2B | Abcam | Cat# ab52484, RRID:AB_1139809 |

| | | |
|---|---|---|
| Mouse monoclonal anti-Histone H2BK120ub | Active Motif | Cat# 39623, RRID:AB_2793279 |
| Mouse monoclonal anti-Histone H4 | Santa Cruz Biotechnology | Cat# sc-25260, RRID:AB_2118623 |
| Mouse monoclonal anti-Vimentin | Developmental Studies Hybridoma Bank | Cat# AMF-17b RRID:AB_528505 |
| Mouse monoclonal anti-Tubulin | Millipore Sigma | Cat# T6199, RRID:AB_477583 |
| Rabbit polyclonal anti-Actin | Abcam | Cat# ab1801, RRID:AB_302617 |
| Mouse monoclonal anti-β-Actin | Sigma-Aldrich | Cat# A5441, RRID:AB_476744 |
| Mouse monoclonal anti-p53 | Santa Cruz Biotechnology | Cat# sc-126 RRID:AB_628082 |
| Mouse monoclonal anti-ETS-1 | Proteintech | Cat# 66598-1-Ig RRID:AB_2881958 |
| Rabbit monoclonal anti-ETS-1 | Cell Signaling Technology | Cat# 14069S RRID:AB_2798383 |
| Rabbit polyclonal anti-IFIT2 | Novus Biologicals | Cat# NBP1-31164 RRID:AB_2121948 |
| Rabbit polyclonal anti-ZAP | Proteintech | Cat# 16820-1-AP RRID:AB_2728733 |
| Rabbit polyclonal anti-STAT1 | Proteintech | Cat# 10144-2-AP RRID:AB_2286875 |
| Mouse monoclonal anti-pSTAT1 | BioLegends | Cat# 686402 RRID:AB_2616883 |
| Mouse monoclonal anti-Vaccinia Virus A27L | Santa Cruz Biotechnology | Cat# sc-58210, RRID:AB_632582 |
| Mouse anti-Vaccinia Virus F4L | (Gammon et al., 2010) | N/A |
| Mouse anti-Vaccinia Virus I3L | (Gammon and Evans, 2009) | N/A |
| Rabbit IgG Isotype Control | Invitrogen | Cat# 10500C, RRID:AB_2532981 |

| | | |
|---|---|---|
| Mouse normal IgG Isotype Control | Santa Cruz Biotechnology | Cat# sc-2025, RRID:AB_737182 |
| Mouse IgG Isotype Control ChIP Grade | BioLegend | Cat# 401506 RRID: N/A |
| Donkey anti-Mouse Secondary, Alexa Fluor 647 | Invitrogen | Cat# A-31571 RRID:AB_162542 |
| Donkey anti-Mouse Secondary, Alexa Fluor 568 | Invitrogen | Cat# A10042 RRID:AB_2757564 |
| Mouse anti Flag M2 Affinity Gel | Sigma | Cat# A2220 RRID:AB_10063035 |
| Virus Strains | | |
| VV-FL-GFP | (Rex *et al.*, 2018) | N/A |
| ∆A51R-FL-GFP | This paper | N/A |
| $\Delta A51R^{FA51R}$ | (Gammon *et al.*, 2014) | N/A |
| ∆A51R | (Gammon *et al.*, 2014) | N/A |
| $\Delta A51R^{FA51R158\text{-}162Ala}$ | This paper | N/A |
| VSV-GFP | (Gammon *et al.*, 2014) | |
| YFV-17D-Venus (YFV-Venus) | (Schoggins *et al.*, 2011) | N/A |
| NEB Stable Competent E.coli (High Efficiency) | NEB | C3040H |
| E. cloni® Chemically Competent Cells | Biosearch Technologies | 60108 |
| BL21(DE3) cells | Agilent | 200131 |
| Chemicals, Peptides, and Recombinant Proteins | | |
| Luciferase Assay Reagent | Promega | E1483 |
| Reporter Lysis 5X Buffer | Promega | E3971 |
| Recombinant H3-H4 | Diagenode | C23010013 |
| Recombinant H2A-H2B | Diagenode | C23010010 |

| | | |
|---|---|---|
| Recombinant ULP-1 SUMO Protease | Sigma | SAE0067-2500UN |
| HisTrap HP 1 ml column | GE Healthcare | 17524701 |
| HiTrap DEAE 5 ml column | GE Healthcare | 17505501 |
| Superdex 200 Increase 10/300 GL column | GE Healthcare | 28990944 |
| Flag Peptide | Sigma-Aldrich | F4799 |
| HIS-Select® Nickel Magnetic Agarose Beads | Sigma-Aldrich | H9914 |
| Tubulin protein (> 99% pure) porcine brain | Cytoskeleton Inc. | T240 |
| GTP | Cytoskeleton Inc. | BST06 |
| Paclitaxel | Sigma | T7402 |
| TGX Stain-Free™ FastCast™ Acrylamide Kit, 10% | Bio-Rad | 1610183 |
| Isopropylthio-β-galactoside (IPTG) | RPI | I560000 |
| Imidazole | Sigma | I2399 |
| Octyl-beta-D-thioglucopyranoside | AFG Scientific | 148922 |
| Dithiothreitol (DTT) | Fisher | BP172 |
| Ethylenediaminetetraacetic acid (EDTA) | Fisher | BP2482 |
| ProLong Diamond Antifade Mountant with DAPI | Invitrogen | P36962 |
| Fibroblast Growth Kit-Low serum | ATCC | PCS-201-041 |
| Fibroblast Basal Medium | ATCC | PCS-201-030 |
| Dulbecco's Modified Eagle's Medium - high glucose | Sigma-Aldrich | D6429 |

| | | |
|---|---|---|
| Minimum Essential Medium Eagle | Sigma-Aldrich | M4655 |
| EX-CELL 420 Serum-Free Medium for Insect Cells | Sigma-Aldrich | 14420C |
| Grace's Insect Medium | Sigma-Aldrich | G8142 |
| Dulbecco's Phosphate Buffered Saline | Sigma-Aldrich | D8662 |
| Antibiotic Antimycotic Solution (100x) | Sigma-Aldrich | A5955 |
| MEM Nonessential Amino Acid Solution | Corning | 25025CI |
| L-glutamine Solution | Corning | 25005CI |
| FB Essence | Avantor Seradigm | 3100 |
| FBS | Atlanta Biologicals | S12450 |
| Opti-MEM Reduced Serum Medium | Gibco | 31985070 |
| Sf-900 II Serum Free Medium | Gibco | 10902096 |
| Lipofectamine 2000 Transfection Reagent | Invitrogen | 11668019 |
| Cellfectin II Reagent | Gibco | 10362100 |
| LI-COR Odyssey Blocking Buffer | LI-COR | 92740010 |
| Econo-Column® Chromatography Columns | Bio-Rad | 7372512 |
| Critical Commercial Assays | | |
| SuperScript™ III One-Step RT-PCR System | Invitrogen | 12574026 |
| MEGAscript™ RNAi Kit | Invitrogen | AM1626 |

| | | |
|---|---|---|
| RNeasy Mini Kit | Qiagen | 74104 |
| SUMO-1 detection kit | Cytoskeleton | BK165 |
| SUMO-2/3 detection kit | Cytoskeleton | BK162 |
| Subcellular Protein Fractionation Kit for Cultured Cells | Thermo Scientific | 78840 |
| ChIP-IT® Express Chromatin Immunoprecipitation Kits | Active Motif | 53009 |
| Detergent-Free Nuclei Isolation Kit | Invent Biotechnologies | NI024 |
| Deposited Data | | |
| RNA-seq raw data | This paper | Accession# GEO: GSE185829 |
| Experimental Models: Primary Cells and Cell Lines | | |
| Human: Primary neonatal human dermal fibroblasts (NHDF) | Lonza | CC-2509 |
| Human: Lung epithelial carcinoma cells (A549) | ATCC | CCL-185 |
| Human: Bone osteosarcoma epithelial cells (U2OS) | ATCC | HTB-96 |
| Human: U2OS-GFP- hSpt16 | This paper | N/A |
| Human: U2OS-GFP-hSpt16$^{\Delta NLS}$ | This paper | N/A |
| Human: U2OS-GFP-hSpt16$^{I554A}$ | This paper | N/A |
| Human: HeLa-Flag-hSpt16 | This paper | N/A |
| Human: Embryonic kidney cells (293T) | ATCC | CRL-3216 |
| Human: Cervical | ATCC | CCL-2 |

| | | |
|---|---|---|
| epithelial cells (HeLa) | | |
| *Cercopithecus aethiops*: Kidney epithelial cells (BSC-40) | ATCC | CRL-2761 |
| *Oryctolagus cuniculus*,: Corneal cells (SIRC) | ATCC | CCL-60 |
| *Rousettus aegyptiacus*: Fetal cell line (R06E) | DSMZ | ACC-756 |
| *Mesocricetus auratus*: Kidney fibroblast cells (BHK-21) | ATCC | CCL-10 |
| *Mus musculus*: Subcutaneous connective tissue cells (L929) | ATCC | CCL-1 |
| *Lymantria dispar*: Ovarian epithelial cells (LD652) | Laboratory of Dr. Basil Arif | N/A |
| *Spodoptera frugiperda*: pupal ovarian tissue cells (Sf21) | Gibco | B82101 |
| Recombinant DNA | | |
| shRNA: MISSION pLKO.1-puro Empty Vector Control Plasmid DNA | Sigma-Aldrich | SHC001 |
| shRNA: MISSION - Spt16 | Sigma-Aldrich | TRCN0000293348 |
| Lentiviral Plasmid: GFP-hSpt16 pReceiver | GeneCopoeia | EX-T0317-Lv103 |
| Lentiviral Plasmid: GFP-hSpt16$^{\Delta NLS}$ pReceiver | This paper | N/A |
| Lentiviral Plasmid: GFP- | This paper | N/A |

| | | |
|---|---|---|
| hSpt16$^{I554A}$ pReceiver | | |
| Lentiviral Plasmid: Flag-hSpt16 pReceiver | This paper | N/A |
| Lentiviral Plasmid: SSRP1 pReceiver | GeneCopoeia | EX-B0096-Lv122 |
| Plasmid: HA-hSpt16-WT pEZ-M06 | GeneCopoeia | CS-T0317-M06 |
| Plasmid: HA-hSpt16$^{\Delta NLS}$ pEZ-M06 | This paper | N/A |
| Plasmid: HA-hSpt16$^{\Delta 515-534}$ pEZ-M06 | This paper | N/A |
| Plasmid: HA-hSpt16$^{\Delta 535-554}$ pEZ-M06 | This paper | N/A |
| Plasmid: HA-hSpt16$^{\Delta 758-893}$ pEZ-M06 | This paper | N/A |
| Plasmid: HA-hSpt16$^{758-893}$ pEZ-M06 | This paper | N/A |
| Plasmid: HA-hSpt16$^{535-554Ala}$ pEZ-M06 | This paper | N/A |
| Plasmid: HA-hSpt16$^{535-541Ala}$ pEZ-M06 | This paper | N/A |
| Plasmid: HA-hSpt16$^{542-548Ala}$ pEZ-M06 | This paper | N/A |
| Plasmid: HA-hSpt16$^{549-554Ala}$ pEZ-M06 | This paper | N/A |
| Plasmid: HA-hSpt16$^{K:A\ 1-199}$ pEZ-M06 | This paper | N/A |
| Plasmid: HA-hSpt16$^{K:A\ 200-399}$ pEZ-M06 | This paper | N/A |
| Plasmid: HA-hSpt16$^{K:A\ 400-456}$ pEZ-M06 | This paper | N/A |
| Plasmid: HA-hSpt16$^{K:A\ 457-629}$ pEZ-M06 | This paper | N/A |
| Plasmid: HA-hSpt16$^{K:A\ 631-830}$ pEZ-M06 | This paper | N/A |

| | | |
|---|---|---|
| Plasmid: HA-hSpt16$^{K:A\ 831-1047}$ pEZ-M06 | This paper | N/A |
| Plasmid: HA-hSpt16$^{K:A\ 400-629}$ pEZ-M06 | This paper | N/A |
| Plasmid: HA-hSpt16$^{K:A\ 400-456+631-830}$ pEZ-M06 | This paper | N/A |
| Plasmid: HA-hSpt16$^{K:A\ 457-629+631-830}$ pEZ-M06 | This paper | N/A |
| Plasmid: HA-hSpt16$^{K:A\ 504/507/513}$ pEZ-M06 | This paper | N/A |
| Plasmid: HA-hSpt16$^{K507A}$ pEZ-M06 | This paper | N/A |
| Plasmid: HA-hSpt16$^{K513A}$ pEZ-M06 | This paper | N/A |
| Plasmid: HA-hSpt16$^{K520A}$ pEZ-M06 | This paper | N/A |
| Plasmid: HA-hSpt16$^{K:A\ 533/534}$ pEZ-M06 | This paper | N/A |
| Plasmid: HA-hSpt16$^{K555A}$ pEZ-M06 | This paper | N/A |
| Plasmid: HA-hSpt16$^{K596A}$ pEZ-M06 | This paper | N/A |
| Plasmid: HA-hSpt16$^{K596/606A}$ pEZ-M06 | This paper | N/A |
| Plasmid: HA-hSpt16$^{K606A}$ pEZ-M06 | This paper | N/A |
| Plasmid: HA-hSpt16$^{FHI549AAA}$ pEZ-M06 | This paper | N/A |
| Plasmid: HA-hSpt16$^{HI550AA}$ pEZ-M06 | This paper | N/A |
| Plasmid: HA-hSpt16$^{I551A}$ pEZ-M06 | This paper | N/A |
| Plasmid: HA-hSpt16$^{TI553AA}$ pEZ-M06 | This paper | N/A |

| | | |
|---|---|---|
| Plasmid: HA-hSpt16$^{I554A}$ pEZ-M06 | This paper | N/A |
| Plasmid: Flag-GFP pcDNA3 | (Gammon *et al.*, 2014) | N/A |
| Plasmid: Flag-VV A51R pcDNA3 | (Gammon *et al.*, 2014) | N/A |
| Plasmid: Gal4-A51R pB66 | This paper | N/A |
| Plasmid: GFP-A51R$^{151\text{-}334}$ C3 | This paper | N/A |
| Plasmid: GFP-A51R$^{201\text{-}334}$ C3 | This paper | N/A |
| Plasmid: GFP-A51R$^{251\text{-}334}$ C3 | This paper | N/A |
| Plasmid: Flag-A51R$^{158\text{-}162Ala}$ pcDNA3 | This paper | N/A |
| Plasmid: Flag-A51R$^{R275A/K295A/K302A}$ pcDNA3 | This paper | N/A |
| Plasmid: pET22b | Sigma | 69744 |
| Plasmid: His-A51R pET22B | This paper | N/A |
| Plasmid: His-A51R$^{158\text{-}162Ala}$ pET22B | This paper | N/A |
| Plasmid: His-A51R$^{R275A/K295A/K302A}$ pET22B | This paper | N/A |
| Plasmid: His-SSRP1 pET22B | This paper | N/A |
| Plasmid: CPXV (strain Brighton) Flag-A51R pcDNA3 | (Gammon *et al.*, 2014) | N/A |
| Plasmid: ECTV (strain Moscow) Flag-A51R pcDNA3 | (Gammon *et al.*, 2014) | N/A |
| Plasmid: YLDV (strain Davis) Flag-A51R pcDNA3 | (Gammon *et al.*, 2014) | N/A |
| Plasmid: MPXV (clinical isolate gp161) Flag-A51R pcDNA3 | This paper | N/A |
| Human siRNA targeting sequences | | |
| siRNA targeting | Sigma-Aldrich | SASI_Hs01_00143917 |

| | | |
|---|---|---|
| sequence: hSpt16-A: ccgugaaggcucccuaguaa | | |
| siRNA targeting sequence: hSpt16-B: caacuggacuaaaaucauga | Sigma-Aldrich | SASI_Hs01_00143919 |
| siRNA targeting sequence: SSRP1-A: cagcuaagcgagagcuucaa | Sigma-Aldrich | SASI_Hs01_00216184 |
| siRNA targeting sequence: SSRP1-B: caccaaaacgaugacgcaga | Sigma-Aldrich | SASI_Hs01_00216186 |
| siRNA targeting sequence: ETS-1-A: cuuuguuggggacaucuuau | Sigma-Aldrich | SASI_Hs01_00173246 |
| siRNA targeting sequence: ETS-1-B: cuggagcuuuucccucccc | Sigma-Aldrich | SASI_Hs01_00173248 |
| siRNA targeting sequence: ETS-1-C: gactacccctcggtcattct | Sigma-Aldrich | SASI_Hs01_00173250 |
| siRNA targeting sequence: ETS-2-A: agucauucaucagcuggacu | Sigma-Aldrich | SASI_Hs01_00130872 |
| siRNA targeting sequence: ETS-2-B: cggcaugaauggccagaugc | Sigma-Aldrich | SASI_Hs01_00130873 |
| siRNA targeting | Sigma-Aldrich | SASI_Hs01_00130879 |

| | | |
|---|---|---|
| sequence: ETS-2-C: tgaatgatttcggaatcaag | | |
| siRNA targeting sequence: HMGA1-A: aagccccatctcatcctggc | Sigma-Aldrich | SASI_Hs01_00134689 |
| siRNA targeting sequence: HMGA1-B: gggggcgccctctctgctcc | Sigma-Aldrich | SASI_Hs01_00134691 |
| siRNA targeting sequence: HMGA1-C: catctcatcctggcacgccc | Sigma-Aldrich | SASI_Hs01_00134693 |
| siRNA targeting sequence: NR0B1-A: cctctacagcttgctcacta | Sigma-Aldrich | SASI_Hs01_00164534 |
| siRNA targeting sequence: NR0B1-B: cccgcggcagggcagcatcc | Sigma-Aldrich | SASI_Hs01_00164535 |
| siRNA targeting sequence: NR0B1-C: acagcatgctgacgagcgca | Sigma-Aldrich | SASI_Hs01_00164536 |
| siRNA targeting sequence: POU5F1-A: tacaggtgcgtgccaccgtg | Sigma-Aldrich | SASI_Hs01_00224603 |
| siRNA targeting sequence: POU5F1-B: cgaggccagggtctctcttt | Sigma-Aldrich | SASI_Hs01_00224604 |
| siRNA targeting | Sigma-Aldrich | SASI_Hs01_00224605 |

| | | |
|---|---|---|
| sequence: POU5F1-C: gcgtgccaccgtgcccagct | | |
| siRNA targeting sequence: RELB-A: gagattgtcgagcccgtgac | Sigma-Aldrich | SASI_Hs01_00103187 |
| siRNA targeting sequence: RELB-B: caagaaatccacaaacacat | Sigma-Aldrich | SASI_Hs01_00103188 |
| siRNA targeting sequence: RELB-C: agagcaaacggcggaagaaa | Sigma-Aldrich | SASI_Hs01_00103189 |
| siRNA targeting sequence: STAT5A-A: tcctgtttgagtctcagttc | Sigma-Aldrich | SASI_Hs01_00086130 |
| siRNA targeting sequence: STAT5A-B: cctgaccatgtactcgatca | Sigma-Aldrich | SASI_Hs01_00086131 |
| siRNA targeting sequence: STAT5A-C: agaggagaagttcacagtcc | Sigma-Aldrich | SASI_Hs01_00086132 |
| siRNA targeting sequence: TRERF1-A: ctctctgtttgaggccaaag | Sigma-Aldrich | SASI_Hs01_00018474 |
| siRNA targeting sequence: TRERF1-B: gacacacacaaggccacact | Sigma-Aldrich | SASI_Hs01_00018475 |
| siRNA targeting | Sigma-Aldrich | SASI_Hs01_00018476 |

| | | |
|---|---|---|
| sequence: TRERF1-C: aatgcacacatgaaaactca | | |
| Oligonucleotides used for ChIP-PCR | | |
| ETS1PCR | FOR: 5’- CCGTCTTGATGATGGTGAGAG<br>REV: 5’- CTTCGCCGCAAAGTGCCAAC | For ChIP-PCR |
| Oligonucleotides used for A51R mutagenesis | | |
| GFP-A51R-Δ1-150 C3 | FOR: 5’-TCAAGCTTATGTTCTTCACCAAGGGCAACGCTTCC<br>REV: 5’-CAGAATTCTTAGCGGGAAGCGATACCGTAGACG | To clone A51R 151-334 into C3 HindIII/EcoRI |
| GFP-A51R-Δ1-200 C3 | FOR: 5’-TCAAGCTTATGAAGTTCTACATCAGCATCAAGTCCGACTACTGC<br>REV: 5’-CAGAATTCTTAGCGGGAAGCGATACCGTAGACG | To clone A51R 201-334 into C3 HindIII/EcoRI |
| GFP-A51R-Δ1-250 C3 | FOR: 5’-TCAAGCTTATGTGCTGCTACTTCGAGCCCCAGATCCGTATCCTG<br>REV: 5’-CAGAATTCTTAGCGGGAAGCGATACCGTAGACG | To clone A51R 251-334 into C3 HindIII/EcoRI |
| GFP-A51R-Δ1-303 C3 | FOR: 5’-TCAAGCTTATGATCGCTCTGTCCGTGGCTGGCAACCTCATCCG<br>REV: 5’-CAGAATTCTTAGCGGGAAGCGATACCGTAGACG | To clone A51R 304-334 into C3 HindIII/EcoRI |
| Oligonucleotides used for hSpt16, SSRP1, and mCherry-His PCR & Mutagenesis | | |
| Flag-hSpt16 pReceiver | FOR: 5' CTAGAGACTACAAGGACCACGACGGCGATTATAAGGATCACGACATCGACTACAAAGACGACGATGACAAGG<br>REV: 5' TCTGATGTTCCTGGTGCTGCCGCTAATATTCCTAGTGCTGTAGCTGATGTTTCTGCTGCTACTGTTCCTTAA | To insert 3XFlag tag in place of eGFP tag (XbaI/EcoRI) upstream of hSpt16 gene |
| HA-hSpt16-K504A pEZ-M06 | FOR: 5’-GGCGAGCAGCAGATCCAGGCTGCCCGCAAGTCCAACGTG<br>REV: 5’-CACGTTGGACTTGCGGGCAGCCTGGATCTGCTGCTCGCC | SDM of K504A hSpt16 in pEZ-M06 |
| HA-hSpt16-K507A pEZ-M06 | FOR: 5’-CAGCAGATCCAGAAGGCCCGCGCATCCAACGTGTCCTATAAGAAC<br>REV: 5’-GTTCTTATAGGACACGTTGGATGCGCGGGCCTTCTGGATCTGCTG | SDM of K507A hSpt16 in pEZ-M06 |
| HA-hSpt16-K513A pEZ-M06 | FOR: 5’-CGCAAGTCCAACGTGTCCTATGCAAACCCCTCCCTGATGCCCAAG<br>REV: 5’-CTTGGGCATCAGGGAGGGGTTTGCATAGGACACGTTGGACTTGCG | SDM of K513A hSpt16 in pEZ-M06 |
| HA-hSpt16-K520A pEZ-M06 | FOR: 5’-GAACCCCTCCCTGATGCCCGCTGAGCCCCATATCCGGGAG<br>REV: 5’-CTCCCGGATATGGGGCTCAGCGGGCATCAGGGAGGGGTTC | SDM of K520A hSpt16 in pEZ-M06 |
| HA-hSpt16-K533/534A pEZ-M06 | FOR: 5’-GGAGATGAAGATCTACATCGACGCGGCGTATGAGACTGTAATCATGCCC | SDM of K533/534A hSpt16 in pEZ-M06 |

| | | |
|---|---|---|
| | REV: 5’-GGGCATGATTACAGTCTCATACGCCGCGTCGATGTAGATCTTCATCTCC | |
| HA-hSpt16-K555A pEZ-M06 | FOR: 5’-GTTCCACATCGCCACCATCGCCAACATTAGCATGTCCGTGG<br>REV: 5’-CCACGGACATGCTAATGTTGGCGATGGTGGCGATGTGGAAC | SDM of K555A hSpt16 in pEZ-M06 |
| HA-hSpt16-K596A pEZ-M06 | FOR: 5’-CCCGAGGCGACTTTCGTCGCTGAGATCACCTACCGGGCC<br>REV: 5’-GGCCCGGTAGGTGATCTCAGCGACGAAAGTCGCCTCGGG | SDM of K596A hSpt16 in pEZ-M06 |
| HA-hSpt16-K696A pEZ-M06 | FOR: 5’-CTTCACCTCCGTCCGGGGCGACGCCGTGGACATCTTGTACAACAAC<br>REV: 5’-GTTGTTGTACAAGATGTCCACGGCGTCGCCCCGGACGGAGGTGAAG | SDM of K696A hSpt16 in pEZ-M06 |
| HA-hSpt16-K606A pEZ-M06 | FOR: 5’-CCGGGCCTCAAACATCGCTGCCCCCGGCGAGCAGAC<br>REV: 5’-GTCTGCTCGCCGGGGGCAGCGATGTTTGAGGCCCGG | SDM of K606A hSpt16 in pEZ-M06 |
| HA-hSpt16-758-893 pEZ-M06 | FOR: 5’-TAGAATTCGGTACCATGCATGACCGGGACGACCTCTATGCCGAGCAGATGGAGCGG<br>REV: 5’-TACTCGAGTCAGTACTTCAGGTCGCAGGAGTTCAACCACTCCTTGATGGG | To clone hSpt16 758-893 into pEZ-M06 EcoRI/XhoI |
| His-SSRP1 pET22b | FOR: 5' -TACATATGCATCATCACCATCATCATGAATTCGAGAACCTTTATTTCCAGGGCGCAGAGACACTGGAGTTCAAC<br>REV: 5'- AGGCGGCCGCTTACTCATCGGATCCTGACGCTG | For PCR/cloning His-SSRP1 into pET22b NdeI/NotI |
| mCherry-His pET22b | FOR: 5'- TGACATGTTGATGGTGAGCAAGGGCGAGGAG<br>REV: 5'-TGGCGGCCGCTTAGTGATGGTGATGGTGATGCTTGTACAGCTCGTCCATGC | For PCR/cloning mCherry-His into pET22b NcoI/NotI |
